# Temporal Variability of Brain-Behavior Relationships in Fine-Scale Dynamics of Edge Time Series

**DOI:** 10.1101/2023.09.02.556040

**Authors:** Sarah A. Cutts, Evgeny J. Chumin, Richard F. Betzel, Olaf Sporns

**Affiliations:** Department of Psychological and Brain Sciences, Indiana University (IU), Bloomington, IN, USA; Program in Neuroscience, IU, Bloomington, IN, USA; Indiana University Network Science Institute, IU, Bloomington, IN, USA; Stark Neurosciences Research Institute, Indiana University School of Medicine, Indianapolis, IN, USA

**Author notes:** Corresponding author: Sarah A. Cutts.

**Keywords:** Time-varying connectivity, brain-behavior associations, functional MRI, resting state

## Abstract

Most work on functional connectivity (FC) in neuroimaging data prefers longer scan sessions or greater subject count to improve reliability of brain-behavior relationships or predictive models. Here, we investigate whether systematically isolating moments in time can improve brain-behavior relationships and outperform full scan data. We perform optimizations using a temporal filtering strategy to identify time points that improve brain-behavior relationships across 58 different behaviors. We analyzed functional brain networks from resting state fMRI data of 352 healthy subjects from the Human Connectome Project. Templates were created to select time points with similar patterns of brain activity. Optimizations were performed to produce templates for each behavior that maximize brain-behavior relationships from reconstructed functional networks. With 10% of scan data, optimized templates of select behavioral measures achieved greater strength of brain-behavior correlations and greater transfer between groups of subjects than full FC across multiple cross validation splits of the dataset. Therefore, selectively filtering time points may allow for development of more targeted FC analyses and increased understanding of how specific moments in time contribute to behavioral prediction.

**Significance Statement:** Individuals exhibit significant variations in brain functional connectivity, and these individual differences relate to variations in behavioral and cognitive measures. Here we show that the strength and similarity of brain-behavior associations across groups vary over time and that these relations can be improved by selecting time points that maximize brain-behavior correlations. By employing an optimization strategy for 58 distinct behavioral variables we find that different behaviors load onto different moments in time. Our work suggests new strategies for revealing brain signatures of behavior.

## Introduction

The link between individual variation in functional connectivity (FC) and behavior is of central interest in human neuroscience, with clear developmental and clinical applications. Such brain-behavior relationships are typically evaluated using FC computed from continuous and extended brain recordings, capturing minutes of time. Longer time frames and multiple scan sessions have been shown to improve reliability of FC [1-4] and support improved ability to distinguish individuals (e.g. ‘fingerprinting’) [5, 6]. However, individual differences in functional networks have been discovered at shorter timescales within a scan [7-9] raising the possibility that brain-behavior associations may be time-dependent. Indeed, extensive literature on time-varying functional connectivity reveals that connectivity patterns fluctuate over time and can be related to different behavioral measures [10-12]. Such approaches typically analyze changes in time series characteristics [10,13-16], transitions or dwell times of derived network states [17-19], instantaneous measures of FC [20,21], or most commonly through smaller sliding windows within a scan session [11,17,22-25]. These fluctuations as well as changes in individual differentiation suggest the possibility that brain-behavior relationships may be stronger during different connectivity patterns only visible from specific moments in time within a scan. Here we assess whether the strength and similarity of brain-behavior relationships between training and testing groups can be improved through targeted selection of time points within a scan session.

Selecting or ‘filtering’ the time series for specific connectivity patterns across subjects could enhance the specificity of behavioral associations. Sporns et al. 2021 show that a resting state scan is composed of the transient appearance of cofluctuations within and between different functional systems. This suggests that apparent connection strength between brain regions changes over time in relation to variations in global coordination. As such, cofluctuations may not always provide information towards a behavior of interest. It is unclear whether data from a full scan includes patterns of cofluctuations that dilute brain-behavior correlations or if separate moments in time better emphasize different behavioral associations. Targeted selection of time points based on specific behavioral or phenotypic queries may provide additional information unseen using the full length of data.

Past work has found evidence of differences between analyses computed on full FC versus targeted selection of frames within a scan. Edge time series (eTS) is one such method that has been used to deconstruct functional connectivity into cofluctuations between brain regions at single time point resolution [27]. With this approach, Esfahlani et al. 2020 found that only a handful of high amplitude moments in time (‘events’) dominate the signal seen in full FC and these moments carry most of the individual differences traditionally studied [29]. Although these ‘events’ contribute the most information towards full FC, they provide only a sparse representation of the whole time series. Although medium to low cofluctuation frames make up the majority of a scan session, their lower amplitude ultimately contributes less towards the averaged pattern of full FC. In this case, targeted selection of time points has already been shown to improve the relation between specific connectivity patterns with behavior as well as reveal additional or unique information from time points with lower representation in full FC. For instance, Cutts et al. 2023 show that subject identifiability [5,30] can be improved above full FC with a discontinuous set of frames making up only 10% of the time series. Other work has found that cognitive function is best predicted by frames distributed across the time series [31]. Additionally, binning time points based on cofluctuation amplitude reveal distinct relations to behavioral prediction [32] as well as clinical phenotype [33] unseen in full FC. These findings suggest that developing techniques to distinguish which moments in time contribute most towards behavioral relationships could aid in improved behavioral and diagnostic prediction.

The examples above filter the time series based on cofluctuation amplitude, but other time series-based metrics or specific connectivity patterns can be uniformly selected across subjects and analyzed. Frames can be filtered based on similarity between empirical nodal activity or connectivity patterns with a predefined template [9]. One approach is through the selection of time points from binarized nodal time series (or ‘bipartitions’ [26]) that are most similar to a predefined binary nodal template. Techniques such as eTS and bipartitions allow computationally expedient alternatives to previous time-varying methods. Simply taking the mean across eTS or the agreement of partition identify across all bipartitioned frames nearly reconstructs full FC [**Fig. 1A,B**; 26]. Accordingly, disparate frames can be selected based on their match with binarized nodal templates and reconstructed into an ‘agreement component’ (AGc) that can be analyzed similarly to FC [**Fig. 1C**; 9, 33]. Access to the temporal deconstruction of the time series also allows for data driven approaches to selectively filter frames for a given study. **Fig. 1D** shows a schematic of an optimization procedure that produces a pattern that can be used to filter frames based on highest similarity to that template. As such, filter templates can be created specifically to isolate frames with given properties of interest.

**Figure 1.**
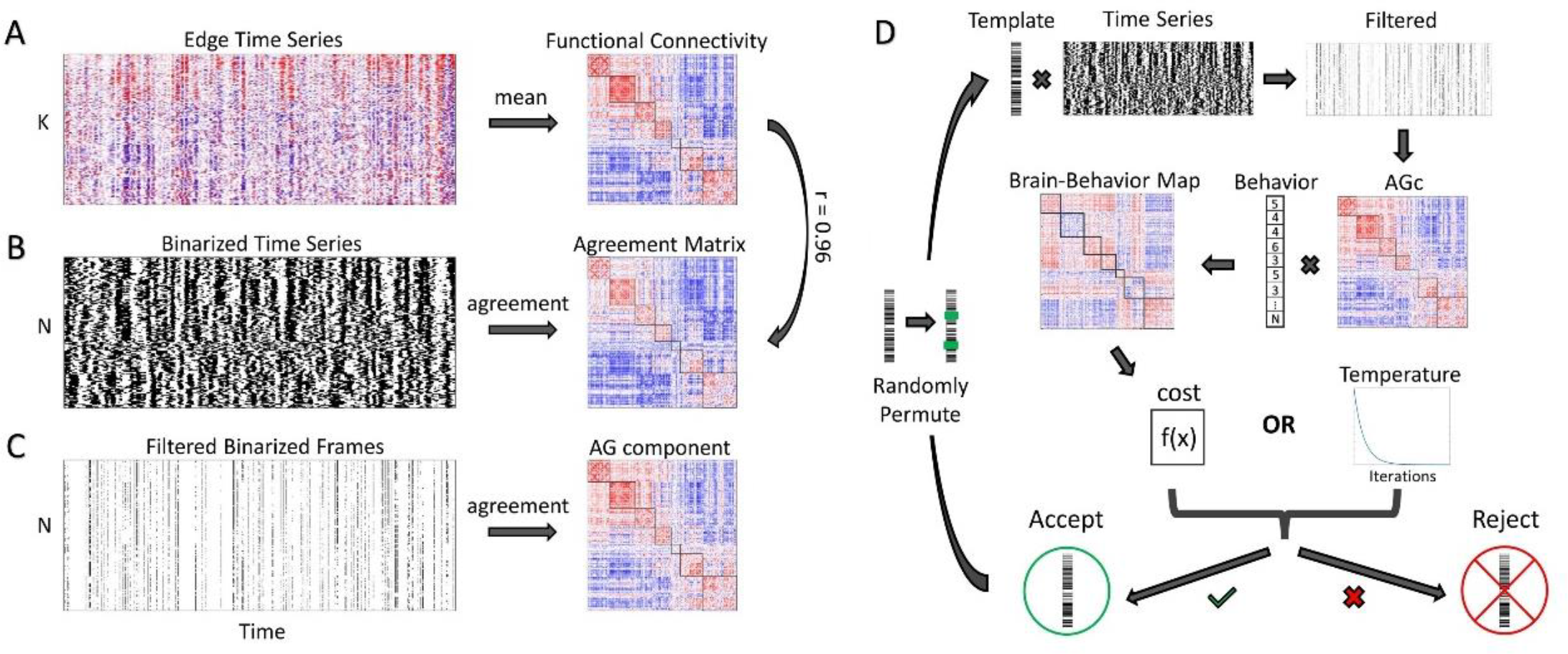
Filtering approach and behavior optimization workflow. The temporal filtering approach is outlined in panels A-C. **(A)** Edge time series (eTS, [27]) forming an edge (K) by time (T) matrix, the mean of which, along T, is the ‘classic’ [N×N] nodal functional connectivity (nFC) matrix. **(B)** Binarized (or bipartitioned) time series are created by thresholding the z-scored time series (TS) at 0 to split each frame into two communities for a matrix of [N×T] bipartitions. The agreement across binarized frames produces a matrix nearly identical to nFC (r = 0.96). **(C)** Frames can be selected (or filtered) from binarized TS and provide a computationally efficient approach for optimizations and machine learning applications. Agreement across filtered frames produces a matrix similar in structure to nFC, here referred as AG components (AGc). **(D)** Optimization pipeline utilizing filtering approach from A-C. The optimization is initialized with a random vector of N length that is adjusted across each of 10,000 iterations to maximize behavioral relations. Normalized mutual information (MI) is used to compare the current template to each time step of the binarized TS. The top 10% (110) time points of T = 1100 with highest MI were selected from the TS. An AGc was created from the selected frames and each edge was correlated to behavioral scores across subjects using Spearman’s ρ. This produced an [NxN] brain-behavior correlation map for the selected template that is translated into a cost function D (here mean of the absolute value of all edges in the behavior correlation map). The cost function is compared to the previous best solution and is either accepted if performance improves or rejected if the ‘temperature’ is high enough to dictate random exclusion of successful results. Between 1-3 nodes of accepted templates are randomly permuted and previous steps are repeated for each iteration.

In this study, we assessed 58 different behaviors in relation to cofluctuation amplitude and developed custom templates to select time points with connectivity patterns that maximize behavioral relationships. Over multiple optimizations, a handful of behaviors revealed that selection of filtered frame sets improved correlation strength and train-test transfer of behavioral associations above full FC. Associated frames were distributed across cofluctuation amplitudes with most selected frames loading onto higher amplitude moments. The few moments in time that overlapped across multiple behaviors displayed greater cofluctuation amplitude and similarity to full FC. This suggests that brain-behavior relationships may be improved if time points are selectively chosen.

## Results

Here, we used filtering to discover the best templates and moments in time that improved brain-behavior correlations. We analyzed 352 subjects from the Human Connectome Project [HCP; 34], selected from the “1200 Subjects” HCP release based on low subject motion and no family relations [35]. Four resting state scans were analyzed from each subject. Fifty-eight different behaviors were selected based on their distribution across cognitive, social, and emotional measures [**Table 1**; 10,36]. First, behaviors were analyzed across cofluctuation amplitudes to assess whether lower amplitude frames with less contribution towards full FC displayed unique behavioral associations. Second, templates were created for each behavior across multiple optimizations using simulated annealing [**Fig. 1D**]. Third, the temporal properties of time points selected from behavior templates were assessed and compared across behavioral measures.

**Table 1.**
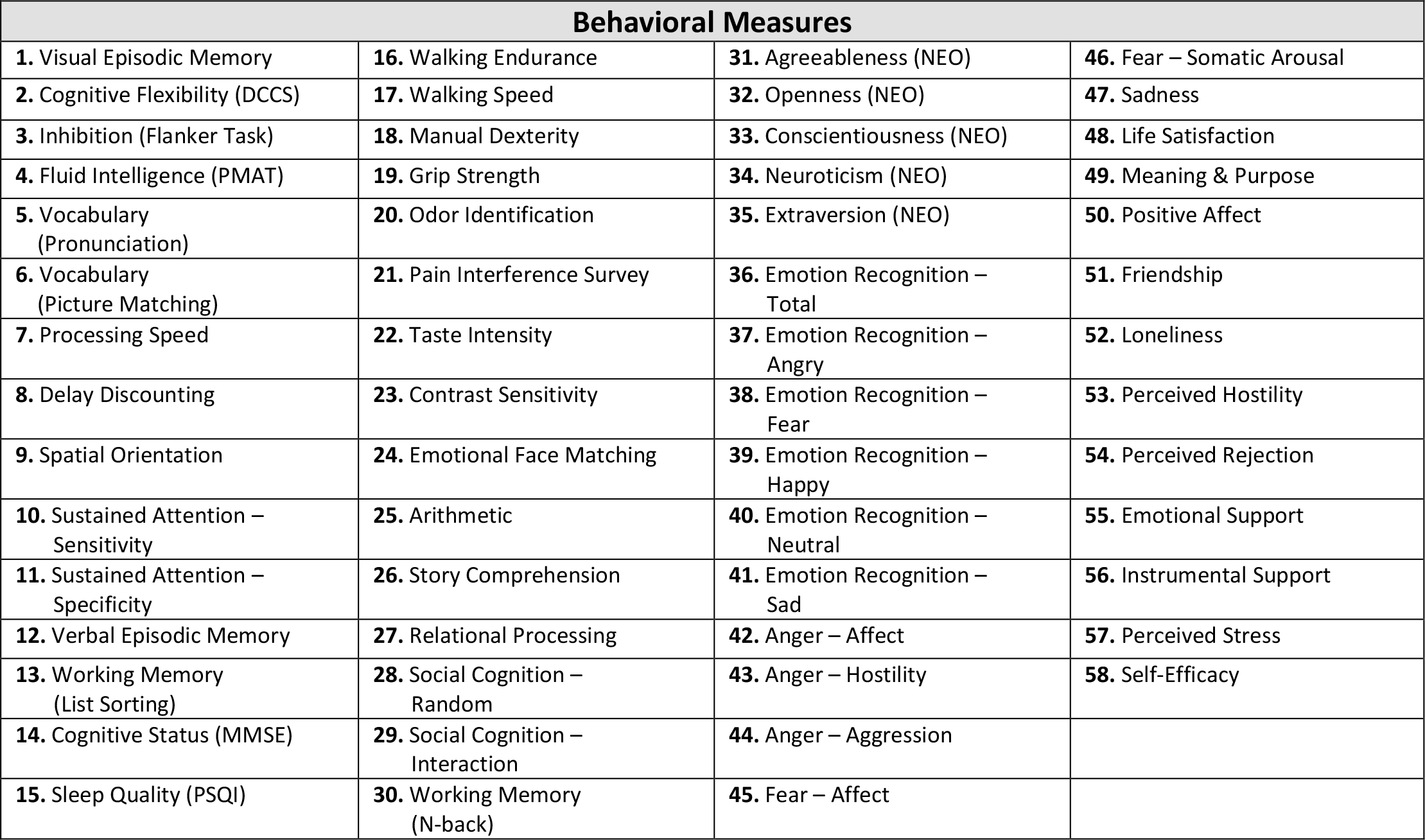
List of 58 behavioral measures. These measures were selected from the Human Connectome Project dataset to span cognitive, social, and emotional measures [10, 36]. Additional information, including field names and descriptions, can be found in the HCP data dictionary [34].

| Behavioral Measures |  |  |  |
| --- | --- | --- | --- |
| 1. Visual Episodic Memory | 16. Walking Endurance | 31. Agreeableness (NEO) | 46. Fear – Somatic Arousal |
| 2. Cognitive Flexibility (DCCS) | 17. Walking Speed | 32. Openness (NEO) | 47. Sadness |
| 3. Inhibition (Flanker Task) | 18. Manual Dexterity | 33. Conscientiousness (NEO) | 48. Life Satisfaction |
| 4. Fluid Intelligence (PMAT) | 19. Grip Strength | 34. Neuroticism (NEO) | 49. Meaning & Purpose |
| 5. Vocabulary (Pronunciation) | 20. Odor Identification | 35. Extraversion (NEO) | 50. Positive Affect |
| 6. Vocabulary (Picture Matching) | 21. Pain Interference Survey | 36. Emotion Recognition – Total | 51. Friendship |
| 7. Processing Speed | 22. Taste Intensity | 37. Emotion Recognition – Angry | 52. Loneliness |
| 8. Delay Discounting | 23. Contrast Sensitivity | 38. Emotion Recognition – Fear | 53. Perceived Hostility |
| 9. Spatial Orientation | 24. Emotional Face Matching | 39. Emotion Recognition – Happy | 54. Perceived Rejection |
| 10. Sustained Attention – Sensitivity | 25. Arithmetic | 40. Emotion Recognition – Neutral | 55. Emotional Support |
| 11. Sustained Attention – Specificity | 26. Story Comprehension | 41. Emotion Recognition – Sad | 56. Instrumental Support |
| 12. Verbal Episodic Memory | 27. Relational Processing | 42. Anger – Affect | 57. Perceived Stress |
| 13. Working Memory (List Sorting) | 28. Social Cognition – Random | 43. Anger – Hostility | 58. Self-Efficacy |
| 14. Cognitive Status (MMSE) | 29. Social Cognition – Interaction | 44. Anger – Aggression |  |
| 15. Sleep Quality (PSQI) | 30. Working Memory (N-back) | 45. Fear – Affect |  |

Cofluctuation amplitude (computed as the root sum square (RSS) across edge cofluctuations at each time step) varies across time. We applied a binning strategy to sort frames into deciles based on RSS strength to assess their relation to behavioral correlations. An AGc was created for each RSS decile bin for every subject and run, and then correlated across all edges with each of the 58 behavioral measures. Strength of brain-behavior correlation was assessed using the mean absolute correlation across all edges. Each behavior consisted of a mean behavioral correlation score for each RSS decile and was compared to 100 AGcs made from randomly selected frames. This provided a general mapping of a behavior’s loading onto RSS amplitude [**Fig. 2**], which were found to cluster behaviors into 5 groups related to RSS profiles [**Fig. 2, *left***; refer to Materials and Methods]. Additionally, a behavior’s loading onto RSS [**Fig. 2, *right***] relates to the strength of behavioral correlations captured in full FC. Here, behaviors that mainly load onto higher amplitude frames (found in the first and second cluster) have a slight increased presence in full FC [**Fig. 2, *middle***] than behaviors that load onto intermediate to low amplitude frames (p = 0.0227, independent sample t-test). This suggests that behaviors that load onto lower amplitude frames may be partially obscured in analyses utilizing full FC.

**Figure 2.**
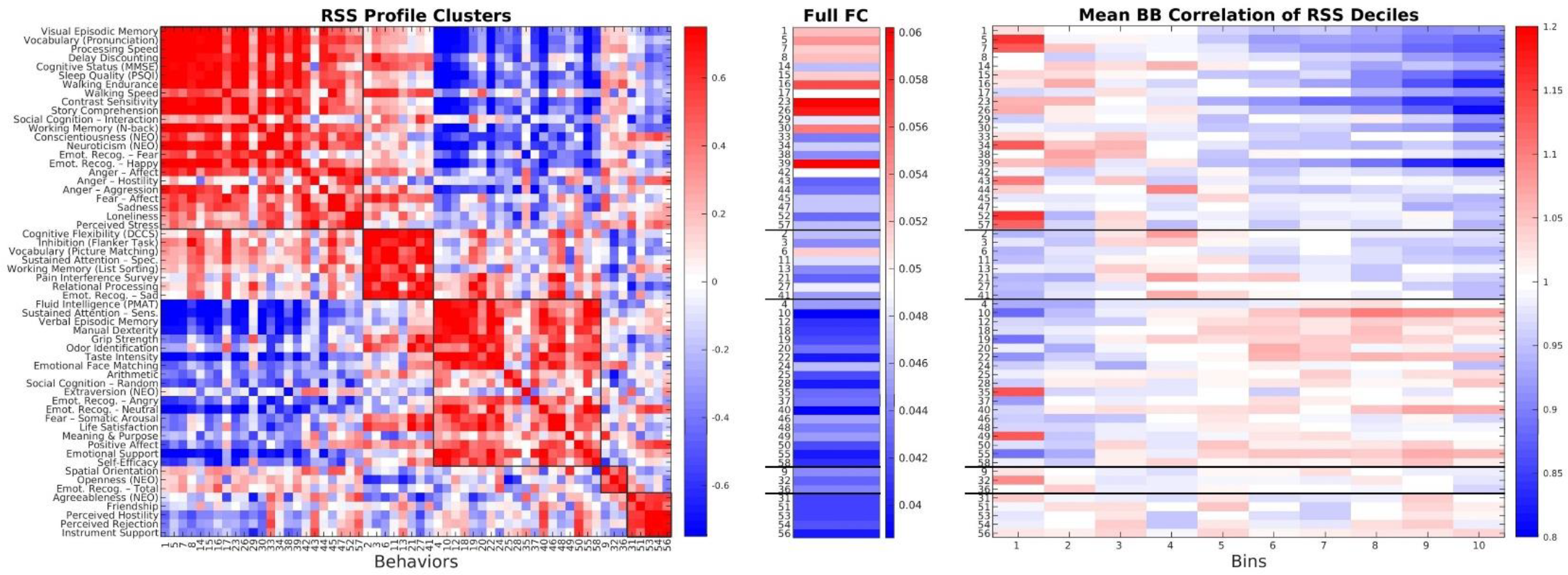
RSS decile binning. (*Left*) Community partition of RSS decile behavioral correlation profiles for 58 behavioral measures from the HCP dataset. A similarity matrix of RSS profiles was clustered into communities using a variant of multiresolution consensus clustering (refer to Methods) to create groups with denser connections (similarities) within communities than between. **(*Middle*)** Mean brain-behavior correlations across all edges as seen in full FC for each behavior and ordered based on derived community identity. **(*Right*)** Vectorized decile bins used in the clustering algorithm (*Left*). The time series was divided into 10 subsets of frames based on RSS amplitude and here average brain-behavior correlations were recorded for each behavior and bin then compared to results from 100 randomly sorted bins.

A data-driven optimization approach was then used to determine which frames systematically carry higher behavioral signal. Binary nodal templates were optimized to provide the strongest behavioral correlations from the top 10% of filtered frames. One-hundred optimizations were performed separately for the 58 behaviors where each iteration created a filtering template from 80% of subjects (282 training set) and analyzed results from the corresponding template on the remaining 20% of individuals (70 testing set). A new randomized training and testing set were used for each of the 100 cross validations. Each optimization was initialized with a random template that was used to filter the top 10% of similar frames from all scans of each subject. Filtered frames were aggregated into an AGc for each subject and used to create an edgewise brain-behavior correlation map. The strength of behavior correlations from this map was used as the cost function to create filtering templates across multiple optimizations. Each step of the optimization accepts or rejects the current template based on the strength of behavior correlations. Accepted templates are permuted at randomly selected nodes and reintroduced to start a new cycle of the optimization. The end result was an optimized template that selects moments in time for which the cost function is maximal. This procedure was performed separately to produce unique optimized templates for each of the 58 different behaviors.

The optimization results from a high performing example behavior (vocabulary pronunciation) are shown in **Fig. 3** (for an additional example behavior, see **Fig. SI1**). **Fig. 3A** shows the overall strength of behavior correlations across each iteration for 100 cross validations of the optimization on the training subjects. As expected, the optimization succeeded in constructing AGcs with much higher behavioral correlations compared to full FC. **Fig. 3B** shows that the optimized behavior templates filter greater behavioral correlations compared to the initial template as well as full FC for the training subjects. Filters applied to held-out test subjects maintained a similar average of the aggregate measure of brain-behavior correlation [**Fig. 3B, *right***], but did show a significant increase in correlation strength from each cross validation over full FC for this given behavior (p < 10^-3^, paired-sample t-test). Notably, the brain-behavior correlation maps are significantly more similar across training and testing subjects. **Fig. 3C** shows the brain-behavior correlation map from the best performing cross validation for full FC, the initial template, and optimized frames. The optimized results reveal a much stronger correspondence between training and testing groups (additional examples in **Fig. SI2, Fig. SI3**). Improved train-test transfer (based on Spearman correlation between brain-behavior maps) can be seen across multiple cross validations and shows significant improvements in similarity across groups (ρ = 0.33) over full FC (ρ = 0.19; p < 10^-15^, paired-sample t-test; **Fig. 3D, *top***). Positive and negative edges from the training set were compared separately to corresponding edges in the testing set. The mean correlation strength of positive edges [**Fig. 3D, *middle***] and negative [**Fig. 3D, *bottom***] were significantly greater than initial templates (p < 10^-15^, paired-sample t-tests) and full FC (p < 10^-13^, paired-sample t-tests). Similar results were found when the number of filtered frames varied [**Fig. SI4, Fig. SI5**]. This shows that filtering frames using optimized templates allowed greater similarity of brain-behavior correlations across training and testing groups as well as improved mean correlation strength for positive and negative edges.

**Figure 3.**
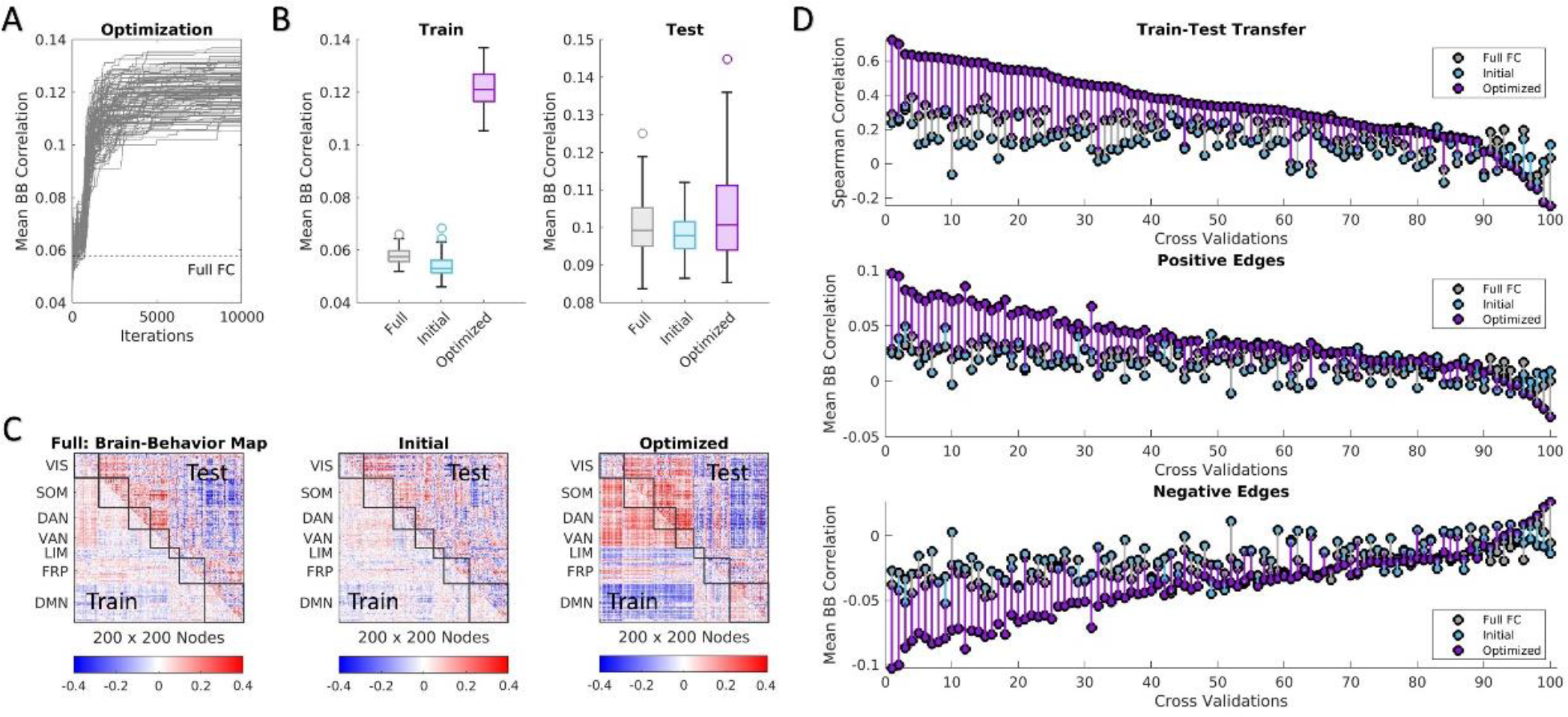
Optimization results for example behavior (vocabulary pronunciation). **(A)** Mean absolute brain-behavior correlations (mean BB) across each iteration of all cross validations compared to full FC. **(B)** Performance in mean BB for brain-behavior correlation maps created from full FC, initial templates, and optimized templates for training (*left*) and testing (*right*) groups across all cross validations. **(C)** Example brain-behavior correlation maps from full FC, initial, and optimized templates from best performing cross validation for training and testing subjects. **(D)** Performance of full FC, initial, and optimized templates assessed for each cross validation using Spearman correlation between training and testing brain-behavior maps (*top*) as well as mean BB of positive edges (*middle*) and negative edges (*bottom*). Positive and negative edge results are shown for the testing group using edge masks selected from brain-behavior correlation maps of training subjects and were ordered based on performance of train-test transfer (*top*).

**Fig. 4** provides a summary of the results across all analyzed behaviors. **Fig. 4A** shows the average mean behavioral correlation from all cross-validations improved for each behavior in the training set. Similar levels of averaged brain-behavior correlations were found across all conditions of the testing set for most behaviors. **Fig. 4B** shows the detailed results as displayed in **Fig. 3**, but for all behaviors. Here, all behaviors with significantly greater train-test transfer of behavioral correlations compared to full FC (p < 0.01, one-tailed t-test uncorrected) also maintained significantly greater strength of positive and negative edges in the testing set. Similar results for p < 0.001 as well as example AGcs and brain-behavior correlation maps across multiple behaviors are displayed in **Fig. SI6** and **Fig. SI7**. A handful of behaviors also showed both greater transfer and correlation strength or only greater correlation strength than full FC. The filtered frames from the optimization were further matched to their corresponding RSS decile bin (shown in **Fig. 2**). **Fig. 4C** relates the count of optimized frames that belong in each RSS bin to the average binning identities of 100 randomly selected bins. Across all cross validations of each behavior, optimized templates select for frames with higher RSS and similarity to FC than the initial template while selecting for similar levels of motion based on framewise displacement [**Fig. 4D**]. However, compared to the highest RSS decile bin, optimized frames maintain lower RSS (p = 0, independent sample t-test) and lower similarity to FC (p < 10^-245^, independent sample t-test) suggesting that filtering recruits frames from lower cofluctuation amplitudes to improve behavioral correlations. These improvements in behavioral correlations and train-test transfer relate to increased variability between subjects. Filtering time points using behavioral templates increased the differentiation of component patterns from full FC [**Fig. SI8, Fig. SI9**] and subject variability [**Fig. SI10**].

**Figure 4.**
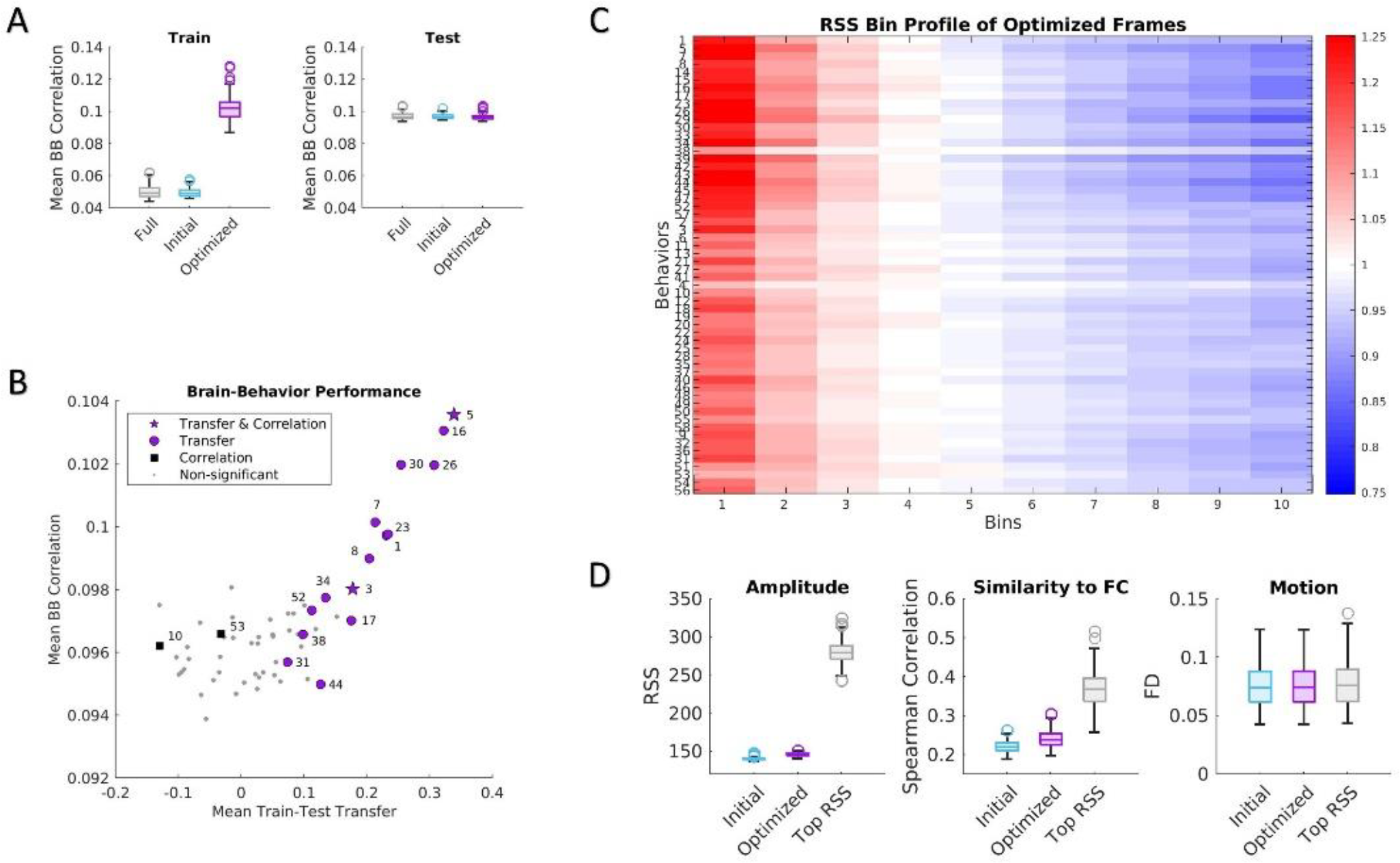
Optimization summary across all behaviors. **(A)** Mean brain-behavior correlations for every behavior using full FC, initial, and optimized templates for training (*left*) and testing (*right*) groups. **(B)** All 58 behaviors plotted as strength of behavior correlations of testing subjects versus train-test transfer (mean Spearman correlation between training and testing group brain-behavior maps). Across all cross validations, each behavior was compared to full FC for train-test transfer, strength of behavior correlation of all edges, and strength of behavior correlations of positive and negative edge sets. Some behaviors showed significant improvements (p < 0.01) over full FC in all mentioned categories (star), improvements in train-test transfer (circle), and mean strength of all correlations (square). Behaviors labeled in purple additionally showed greater strength of mean correlations both in positive and negative edges. **(C)** Frequency of optimized frames assigned to each RSS amplitude decile bin, ordered from highest RSS to lowest. Behaviors ordered based on RSS bin clusters from Figure 2. **(D)** Mean characteristics of initial and optimized frames across all behaviors for RSS (*left*), similarity to full FC (*middle*), and motion (framewise displacement; *right*).

So far, the results have been aggregated over the selected frames of all subjects and runs as well as multiple cross validations, but the optimization pulls frames from specific time points whose properties can be analyzed directly. Out of the random 80/20 (train/test) splits of the 100 cross validations, subjects were selected multiple times across both groups. Over multiple cross validations of an individual subject, the optimizations consistently targeted a select set of specific time frames. **Fig. 5A (*top*)** shows an example heatmap for the frequency of frame selection for the example behavior (vocabulary pronunciation) across 7 cross validations of testing set results of a select subject and run. Each column of the matrix depicted below [**Fig. 5A (*middle*)**] consists of the corresponding heatmaps of each subject and run. Heatmaps of all subjects and runs were compared to 10,000 circular shifted nulls to determine a significance score for each frame (refer to Materials and Methods). Frames were then selected based on a threshold of p < 0.01 (uncorrected) for significantly greater temporal consistency across optimizations than chance. A stricter threshold of p < 0.001 provides similar results (shown in **Fig. SI11** and a separate behavior in **Fig. SI12**). The number of frames found to be significantly consistent among cross validations was much higher for the optimized templates [**Fig. 5B**; mean = 29.02] than for the initial template used at the start of the optimization [**Fig. 5A**; mean = 3.30; p = 0, paired-sample t-test]. This suggests that the optimized templates allowed for the consistent selection of time points in held-out test data despite multiple independent initializations of the optimization with different groupings of subjects.

**Figure 5.**
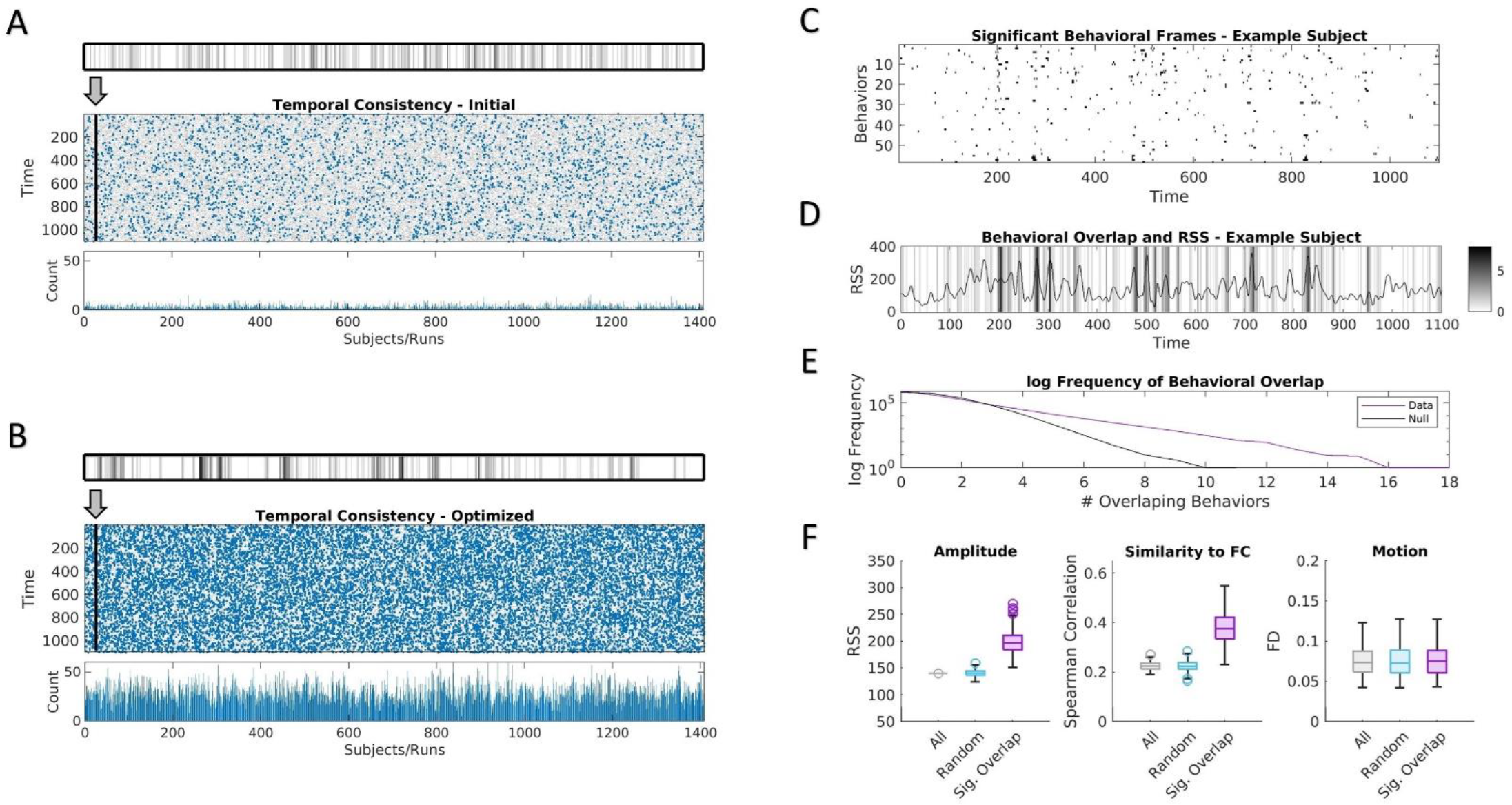
Temporal consistency and behavioral overlap. **(A)** Temporal consistency of frames filtered using the initial template of example behavior (vocabulary pronunciation) across multiple cross validations. Results are shown for cross validation training templates applied to subjects in the testing groups. Frequency of frames selected across all considered cross validations for each time point are displayed for every subject and run. Example subject shown (*top*) as plotted for every subject and run (*middle*). Time points with significant consistency between cross validations (p < 0.01; compared to 10,000 circshifted nulls) plotted in blue and the frequency of occurrence of these significant time points are shown (*bottom*). **(B)** Temporal consistency of optimized templates of example behavior (vocabulary pronunciation). **(C)** Frames with significant consistency as shown in B plotted for each behavior across time for one example subject and run. **(D)** Overlap across behaviors of significant time points as in C for the example subject and run. Corresponding RSS values displayed. **(E)** Plot of log frequency of instances of behavioral overlap of significant frames across all subjects and runs of the test group compared to distribution of 10,000 circshifted nulls. **(F)** Mean characteristics of frames across all subjects and runs for all frames, matching number of randomly selected frames, significant behavioral overlap (p < 0.01) for RSS (*left*), similarity to FC (*middle*), and motion (framewise displacement; *right*).

These significant frames were then compared across behaviors. In **Fig. 5C**, the significant frames are shown for each behavior of an example subject and run. From that, selected frames across behaviors appear to largely occur independently of one another but a handful of frames reveal overlap across behaviors. **Fig. 5D** shows the behavioral overlap of the example subject and run shown in **Fig. 5C** and compares the frame selection to the RSS amplitude time series. Across all subjects and runs, there were time points with greater behavioral overlap compared to 10,000 circular shifted nulls [**Fig. 5E**]. The time points with significant behavioral overlap (p < 0.01, uncorrected) were averaged across all runs of each subject and general framewise properties were compared to results from initial and all frames. In **Fig.5F**, moments in time with significant behavioral overlap display increased RSS amplitude, higher similarity to full FC, and non-significant differences in head motion compared to initial templates and all frames.

## Discussion

Functional connectivity is becoming increasingly used towards the development of brain-based biomarkers of healthy cognition, demographics, and clinical diagnostics [33, 37-46]. There are already several machine learning and predictive models that have been applied towards improving behavioral prediction and developing a deeper understanding of what networks can tell us about cognition and clinical phenotype [5, 31-32, 36, 47-55]. A large portion of FC research has been done on fMRI data, so methodological constraints and the slow hemodynamic response of the BOLD signal [56-59] has led to the preference of conducting analyses on full scan sessions of several minutes to improve reliability [1-3]. Recent studies, however, suggest that additional behavioral information can be obtained from the analysis of temporal fluctuations 7-12, 60-62]. There are already several approaches within the time-varying literature 10-11, 13-25], but there are few answers regarding how best to analyze connectivity data for specific behaviors or phenotypes of interest. Here, we assess whether filtering frames based on specific patterns of activity (bipartitions) can improve behavioral associations. We used optimizations to develop templates to isolate time points consistently across subjects with stronger behavioral associations and determine whether separate moments are better for different behaviors. We also show that the time points filtered for these behaviors are consistent across multiple optimizations and create transferable brain-behavior associations in held-out subjects. We found that a few moments in time improve behavioral associations across multiple behaviors but that some behavioral information loads onto moments that are less prominent in full FC.

Our approach provides a framework to select behaviorally relevant moments in time from a typical scan session consistently across individuals. These results suggest that moments in time carry different signatures of information that are relevant towards specific brain-behavior associations. We found that there are some moments that are more informationally rich than others and that behavioral relations differ based on cofluctuation amplitude as well as ability to be detected from full FC. Moments that enhance behavioral associations differ across behaviors suggesting that filtering can enhance connectivity patterns based on the phenotype or behavior of interest. This is done with resting-state where normally large amounts of data are needed to ensure the researcher is analyzing stable FC across individuals. Rather than analyzing summaries over longer periods of time, finding moments that are similar across subjects in resting-state could fulfill this need with less data and clearer results with the added benefit of improved temporal localization of behaviorally related signatures. Recent work suggests that thousands of subjects are needed to establish strong and reliable brain-behavior associations [63-66]. However, filtering time points based on specific behaviors or phenotypes provides an additional avenue for exploring intersubject variability in behavior. Longer scans would still be required to identify time points that better relate to behavior, but by sorting frames based on their behavioral relevance, more nuanced information may be achieved over otherwise static full FC.

Linking behavioral and phenotypic associations in FC back to mechanistic or neurobiologically plausible sources will likely require improvements in temporal specificity. Along with fluctuations in time-varying FC [11, 13-25] individually distinctive signatures also vary over time 7-9, 32]. As such the relation between intersubject variability in FC and behavioral measures will change based on the connectivity pattern compared across subjects. With enough data, FC stabilizes within subjects [2, 67-68] to provide a clear baseline for intersubject comparisons. However, moments in time that emphasize connections between specific brain regions or functional systems can also provide a scaffold for comparisons that can be related back to intersubject variability of specific behaviors. The origin of these temporal fluctuations in FC remains unclear, with some studies suggesting an origin in neuronal processes [69-70] or network dynamics [71-72] while others interpret FC states as expected stochastic variations around a central tendency that can be observed at the full scan length [73-74]. Regardless of the origin, finding the moments in time that contribute most towards behavioral associations allows for closer inspection of the relation with other neurobiological signatures or noise. Although behaviorally filtered frames from this study did not show systematic differences attributable towards noise (such as head motion, [**Fig. 4D, Fig. 5F**]) and potential confounding variables such as age, sex, and BMI were regressed out, the time points selected for improvements in behavioral correlations could be related to additional physiological or behaviorally relevant time stamps. Some of these improvements in behavioral associations may be driven by variation in physiological signatures typically considered as artifact, but it is important to determine both where such improvements in behavioral associations arise as well as the relationship between potential sources of artifact and their correlations with behavioral or phenotypic traits [54, 75]. Additionally, future work will involve optimizing behavioral relations to movie and task data where the exact stimuli that drive behavior relations can be assessed through comparison to the contents of the movie or task of interest.

This study provides proof of concept for the selective filtering approach and shows that behaviors load onto disparate time points scattered throughout the scan session. The templates created here were optimized on 58 behaviors selected to represent a larger set [34, 76]. Although this provides an overview of different characteristics of behavioral measures in FC, templates for each of these behaviors were produced from the same optimization parameters. Each of these behaviors would benefit from specialized attention to improve targeted frame selection. For instance, we know that behaviors occur at varying timescales, therefore optimization procedures would likely benefit from considering the number of selected frames, sequences of frames, or the use of specific nodes or functional networks. Here, analyzed components were created from 10% of the time series and it remains unclear whether behaviors would benefit from varying the quantity of considered frames (results displayed for an example behavior using 5% [**Fig. SI4**] and 15% frames [**Fig. SI5**]). It is also likely that only part of the template is relevant for isolating behavioral relations of interest and that the rest of the template shows random relations with the time frames in question. This makes a direct interpretation of the templates challenging regardless of their utility in retrieving desired framesets. Lastly, the optimization occasionally produces an exact inversion of the brain-behavior map found in most other cross validations (as seen in the cross validations with worse train-test transfer in **Fig. 3D, Fig. SI3**). This likely occurs when the randomly selected subjects in the testing set involve outliers of the overall trend. A similar issue with the optimization is that for most behaviors a handful of cross validations do no better than the initialized template with low train-test transfer as well as clear lack of functionally relevant structure in edgewise correlations of the brain-behavior map. However, it should be noted that the same cross validations that experience inversions or fail to optimize clear edgewise correlations, also mirror failures or inversions in full FC [**Fig. SI3**], likely relating issues back to the group of subjects analyzed. Overall, successful optimizations repeatedly return the same brain-behavior correlation map as well as filtered time indices across multiple iterations. Additional work is necessary to determine how these templates lock onto and improve behavioral associations.

In summary, we show that behavioral relationships vary across time and do not always occur at the same moments for different behaviors. These behavioral relations map onto specific activity (bipartition) templates that can be used to filter similar functional connectivity patterns across subjects even during resting-state. These optimized behavioral moments occur across RSS amplitudes and therefore may benefit from filtering approaches to emphasize brain-behavior relations not as visible in full FC. Future applications abound, including studies probing for individual differences that involve developmental or clinical cohorts, and that leverage emerging machine learning and A.I. technology.

## Materials and Methods

### Dataset and Acquisition

Analyses were done on data from the Human Connectome Project [HCP; 34]. From the 1200 data release, 352 subjects (54% female, mean age = 29.11+/−3.67, age range = 22–36) selected based on low subject motion, high level of data quality/completeness, low incidence of artifact removal, and no family relations [35]. All procedures and study protocols were approved by the Washington University Institutional Review Board and informed consent was obtained from all participants. Participants were instructed to fixate on a cross during the scan session. Data were collected using a 3T Siemens Connectome Skyra with a 32-channel head coil. Four 15-min resting-state fMRI scans were collected from each participant over 2 days with a gradient-echo EPI sequence, TR = 720 ms, TE = 33.1 ms, flip angle = 52, 2-mm isotropic voxel resolution, and multiband factor = 8. After removal of frames from both the start and the end of the scan session, 1100 time points remained for a scan duration of 14:33 min. Data acquisition and the minimal preprocessing pipeline is described in detail in Glasser et al. 2013.

A selection of 58 behavioral measures were analyzed from a larger set from the 1200 HCP subject release [34, 76] based on their varied distribution across cognitive, social, and emotional measures [10, 36]. The confounds of age, sex at birth, body mass index (BMI), framewise displacement (FD), and FreeSurfer intracranial volume were regressed separately for each cross-validation training and testing group. One-hundred cross validations were created in 80/20 subject splits, where each set consisted of 282 randomly selected training subjects and results were analyzed in the remaining 70 testing subjects.

### Preprocessing

Nilearn signal.clean was used to linearly de-trend, band-pass filter (0.008-0.08 Hz) [78], confound regress, and standardize all functional data. The HCP functional data were cleaned of signal artifacts with ICA-FIX [79]. The functional data were then nuisance regressed using the global signal, the signal derivative, and the squares of each term.

Regional time series were obtained using the Schaefer 200 functional parcellation as provided in the fs_LR surface space [80]. Following nuisance regression, functional data were averaged within each regional node per frame to create 200 spatially distinct time series. Each of the 200 nodes were matched to a set of canonical functional networks as described by Yeo et al. 2011: visual (VIS), somatomotor (SOM), dorsal attention (DAN), ventral attention (VAN), limbic (LIM), frontoparietal (FP), and default mode (DMN).

### Functional Connectivity and Edge Time Series

In this study, functional connectivity (FC) is defined as the statistical dependencies between BOLD time series of every pair of brain regions. Pearson correlation is most commonly used to establish the distance between nodal time series and can be calculated as

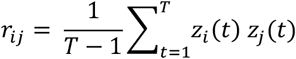

where T defines the total number of time points (*t*) and *z*_*i*_ represents the z-scored value of region *i*. Computing this for every pair of regions produces an [*N*×*N*] matrix with *N* number of nodes. All analyses in FC and on connectivity components were vectorized to include 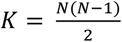unique edges.

FC can be deconstructed into a [*K*×*T*] matrix of all edges (*K*) by time (*T*). This method is referred to as edge time series (eTS) and produces cofluctuation time series between each pair of nodes [27]. Essentially, eTS removes the final averaging step from the Pearson correlation, revealing the instantaneous relationships between the activity of node pairs as

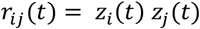

where *r*_*ij*_(*t*) provides a measure of cofluctuation of regions *i*and *j*at time *t*. Averaging eTS across time returns standard nodal functional connectivity (nFC) and therefore can be used to isolate the exact contribution of temporal features that produce FC. eTS was computed for each subject to derive root sum square (RSS) amplitude values at each timepoint as

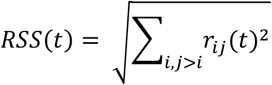

where each time (*t*) is assigned a value related to strength of global cofluctuation across all edges (*K*).

### Bipartitions and Connectivity Components

For each subject and run, the z-scored BOLD time series was binarized by thresholding at zero (BOLD > 0 = 1 and BOLD < 0 = 0). This removed amplitude information while retaining the sign of the signal. An agreement matrix or co-classification matrix [82-83] provides an account of the frequency in which two nodes are assigned to the same community. This is compared to a null model of the expected frequency of community identity given random permutations of nodes while keeping the number and size of clusters fixed. The null was computed as

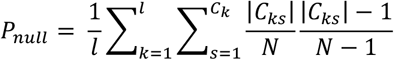

where *l* is the number of samples, *C*_*k*_ the number of clusters in the k-th sample, |*C*_*ks*_ | is the number of nodes in a given cluster of *C*_*k*_, and *N* is the total number of nodes. Here each time point (*t*) was considered as a sample (*k*) and each bipartition group as a cluster (*s*). The null was subtracted from the agreement matrix prior to further analysis. Computing an agreement matrix across the bipartitioned time series produces an [*N*×*N*]matrix across the selected subset of frames that is nearly identical to standard nFC [26]. AGc were created by computing an agreement matrix from subsets of frames within a scan session.

### Amplitude Binning and Frame Filtering

Connectivity components allow for network-based analysis on non-continuous moments in time that reflect exact temporal contributions towards nFC. Frames can be selectively filtered across time based on multiple criteria of interest (refer to [9]). Two filtering methods were deployed. 1) Binning of frames based on a framewise metric. 2) Similarity of frames to a preselected template.

Binning can be performed using any desired criteria of interest that temporally matches the functional time series. Frames can then be sorted into bins of an experimentally chosen size based on the amplitude of the framewise metric. Here, frames were sorted into deciles of RSS amplitude to directly test the appearance of brain-behavior relationships from frames which contribute less towards full FC [28].

Template filtering can be performed by computing the similarity between a template and each frame of the time series. Here, distance was computed using the mutual information (MI) between a binarized template (vector of nodal-length) and the bipartitioned nodal time series for each time point. MI was computed as

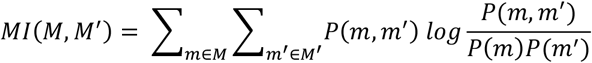

where *m* and *m*^′^ represent modules within the larger partitions of *M* and *M*^′^ under comparison. *P*(*m, m*^′^) gives the fraction of nodes in *m* and *m*^′^ that are present in both modules [84]. We normalized the measure to account for entropies of both *M* and *M*^′^ and rescale the unit interval. The top 10% of frames with the highest normalized MI were selected for connectivity components. Further information on template filtering can be found [9, 26].

### Multiresolution Consensus Clustering

The mean brain-behavior correlations found for each RSS decile were clustered into groups of behaviors with similar profiles. A distance metric (here Spearman correlation) was calculated across all pairs of behaviors using their RSS correlation profiles to produce a [behavior x behavior] matrix that was then clustered. Briefly, modularity maximization is a common approach used to compute data-driven communities defined by denser internal connections than would be expected by chance and weaker connections across communities [85-86]. The modularity quality function is computed as

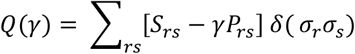

where *S*_*ij*_is the similarity between behavior profiles *r* and *s, P*_*rs*_ is the expected similarity by chance, and *δ*(*σ*_*r*_*σ*_*s*_) is the Kronecker delta function that is equal to 1 when community assignments of *r* and *s*, denoted as *σ*_*r*_ and *σ*_*s*_, are identical and 0 otherwise. The resolution parameter *γ* controls the size of communities that modularity detects. The Louvain algorithm [87] was used to maximize the value of Q, as a representation of the quality of community partition, for each entered value of the resolution parameter *γ*. Here, we used the Potts null model [88] and the full FC matrix, including negative edges. We conduct a general “sweep” over the *γ* parameter by sampling at 1,000 different values, followed by a finer sampling within the range where two-community structure appears (10,000 samples). This creates a set of multiresolution communities [82] which are then transformed into a probabilistic agreement matrix showing the fraction of instances in which nodes *r* and *s* were assigned to the same communities.

### Behavior Optimization

An optimization algorithm was used to create custom templates for each behavior that would isolate framesets which maximized brain-behavior correlations across functional connectivity edges. Due to the similarity between the agreement matrix of binarized BOLD time series and full amplitude FC, both binary templates and AG components were utilized for computational efficiency. The algorithm was initiated with randomized binary vectors of length N (number of nodes). Normalized mutual information was used to find the similarity between these initial templates and the binarized nodal activity of each time point. The top percentage of best matching frames (in this case 10%) were used to produce AG components for all subjects and runs. Brain-behavior correlations were found for each edge of the average AG component across subjects. In each new iteration, the performance of a template was compared to the template of the previous iteration in order to maximize brain-behavior correlations. If the new template outperformed the previous, it was kept and modified for the next iteration. This process was repeated over 10,000 iterations to allow the algorithm to converge on a stable end state.

With this method, there are 2^N^ template permutations which is too large for exhaustive analysis. Therefore, an implementation of the Metropolis-Hastings algorithm [89] was used to efficiently probe the search space. An objective function D was defined as *mean*(*abs*(*BB*)), where *BB* represents the brain-behavior correlations for all edges. Improvement upon this objective function was used in conjunction with simulated annealing [90] to determine the acceptance or rejection of each template iteration. Upon each iteration, a temporary template was created by randomly inverting the binary identities of a handful of nodes from the previous template. The number of altered nodes followed a normal distribution where 1, 2, or 3 nodes were changed with a frequency of 0.68, 0.27 and 0.04, respectively. The success of the temporary template was governed by either the improvement of the objective function or the simulated ‘temperature’ of the system. A cooling schedule was defined as *Temp* = *T*_0_*T*_*exp*_^*h*^, where *T*_0_ defines the initial temperature, *T*_*exp*_ the steepness of the gradient decay, and *h* the iteration number. An annealing criterion of *e*^−Δ*D*/*Temp*^ > *R*(0,1) was used to determine acceptance of the current template, where *Temp* refers to the simulated ‘temperature’ governed by the cooling schedule and *R*(0,1) is a random number between [0,1]. This annealing criterion occasionally rejects local optima in favor of continued exploration earlier in the optimization process. As the ‘temperature’ decays based on the cooling schedule, the algorithm increasingly accepts optimal solutions as governed by the objective function. One set of parameters was selected for all behaviors to ensure stable convergence across repeated runs of the optimization within a reasonable time.

Optimizations were performed across 100 cross validations of 80/20 (train/test) groups for each of the 58 behaviors to evaluate consistency across template outputs. The selected framesets were analyzed for temporal consistency across 7 cross validations for each subject (based on the minimum from the randomized creation of groups). Selected framesets were additionally compared to framewise properties of the time series such as RSS, similarity to full FC, and framewise displacement (FD). Generalizability was tested by applying the optimized templates to the held-out testing subjects for each cross validation and analyzing the similarity between brain-behavior correlation maps of the training and testing sets.

### Temporal Analysis

The temporal location of best matching frames for each optimized behavioral template was recorded for each subject and run. This information was used to compare the consistency of iterations of the optimization as well as determine the overlap between time points retrieved from separate behaviors. The consistency across iterations of the optimization were compared to circshifted nulls (using the MATLAB circshift function) of the selected frame indices for each behavior.

Each iteration of the optimization returned 110 best matching frames regardless of overall fit. Plotted as a binary raster, successful convergence of the optimization is expected to display consistent time frames selected per optimization run. The occurrence of these frame selections were summed across the test group results from multiple cross validations at each moment in time and the frequency of overlap were then assessed for significance against the same number of randomly shifted framesets. The null model was developed by taking the true framesets and randomly shifting them either forwards or backwards in time separately for each cross validation. Frames shifted beyond the boundaries were wrapped around to the opposite side of the time course. This procedure preserved the autocorrelation and temporal spacing between frames while varying the amount of overlap between cross validations. Ten thousand iterations of the null were performed to establish significance in overlap across optimization runs.

The significant timepoints established from this method were recorded for every behavior, subject, and run. Overlap across behaviors was additionally established by comparing significant moments in time across behaviors separately for each subject and run to 10,000 rounds of circshifted frames. The number of instances of behavioral overlap were tested against the null across all subjects to establish the degree to which behaviors load onto similar or disparate time points. Considering that RSS has been found to drive the signal present in full FC [28-29], behavioral overlap was also compared to cofluctuation amplitude.

## Author contributions

S.A.C. and O.S. designed research; S.A.C. performed research and analyzed data; O.S. supervised project; O.S., E.J.C., and R.F.B. edited manuscript; S.A.C. wrote the paper.

## Acknowledgments

This research was supported in-part by NSF grant no. 2023985 (R.B. and O.S.), the National Institute on Aging Training Grant on Alzheimer’s Disease and ADRD (AG071444; E.J.C), and the Indiana University Network Science Institute. This research was supported, in part, by the Lilly Endowment, through its support for the Indiana University Pervasive Technology Institute and, in part, by the Indiana METACyt Initiative [91]. The Indiana METACyt Initiative at Indiana University was also supported, in part, by the Lilly Endowment. Data were provided, in part, by the Human Connectome Project, WU-Minn Consortium (principal investigators: D. Van Essen and K. Ugurbil; 1U54MH091657), funded by the 16 National Institutes of Health (NIH) institutes and centers that support the NIH Blueprint for Neuroscience Research and by the McDonnell Center for Systems Neuroscience at Washington University.

---

**Figure SI1.**
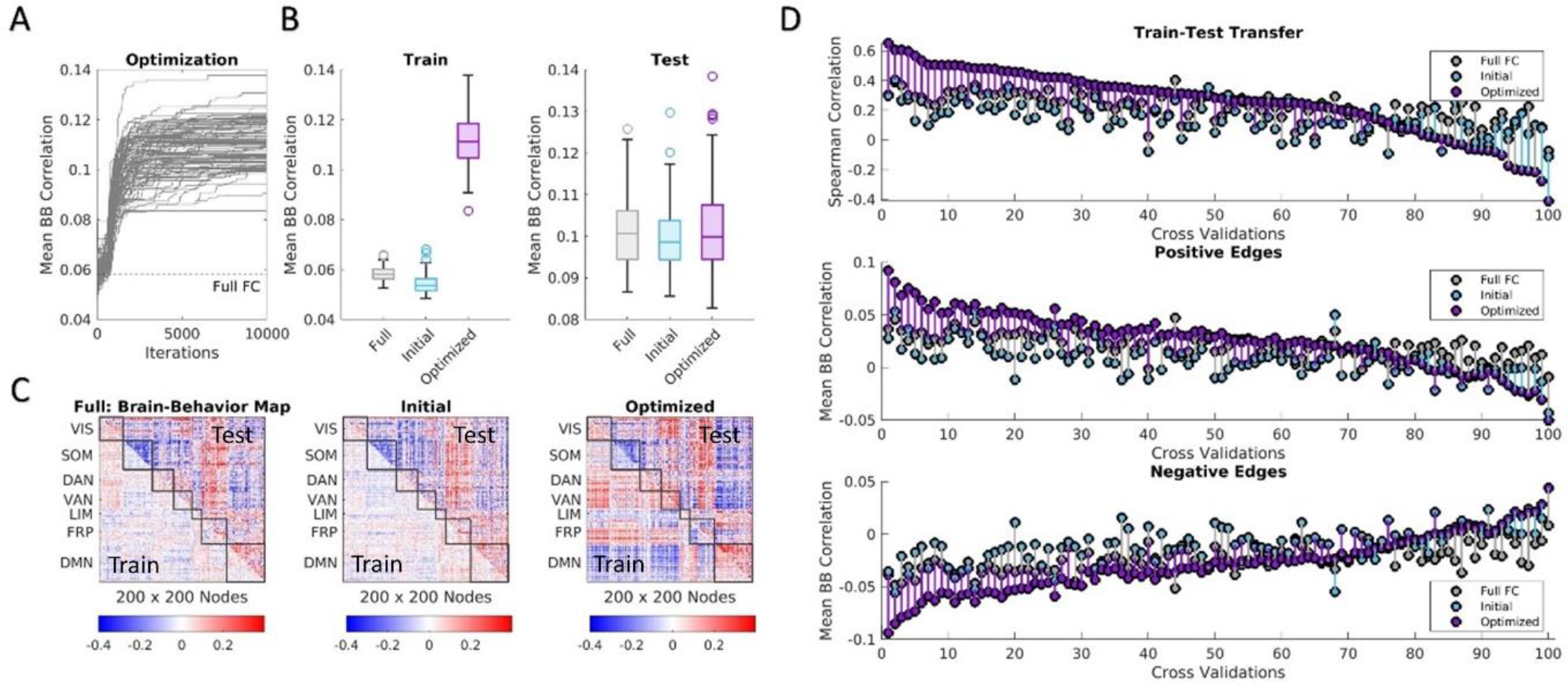
Optimization results for example behavior (working memory). **(A)** Mean absolute brain-behavior correlations (mean BB) across each iteration of all cross validations compared to full FC. **(B)** Performance in mean BB for brain-behavior correlation maps created from full FC, initial templates, and optimized templates for training (*left*) and testing (*right*) groups across all cross validations. **(C)** Example brain-behavior correlation maps from full FC, initial, and optimized templates from best performing cross validation for training and testing subjects. **(D)** Performance of full FC, initial, and optimized templates assessed for each cross validation using Spearman correlation between training and testing brain-behavior maps (*top*) as well as mean BB of positive edges (*middle*) and negative edges (*bottom*). Positive and negative edge results are shown for the testing group using edge masks selected from brain-behavior correlation maps of training subjects and were ordered based on performance of train-test transfer (*top*).

**Figure SI2.**
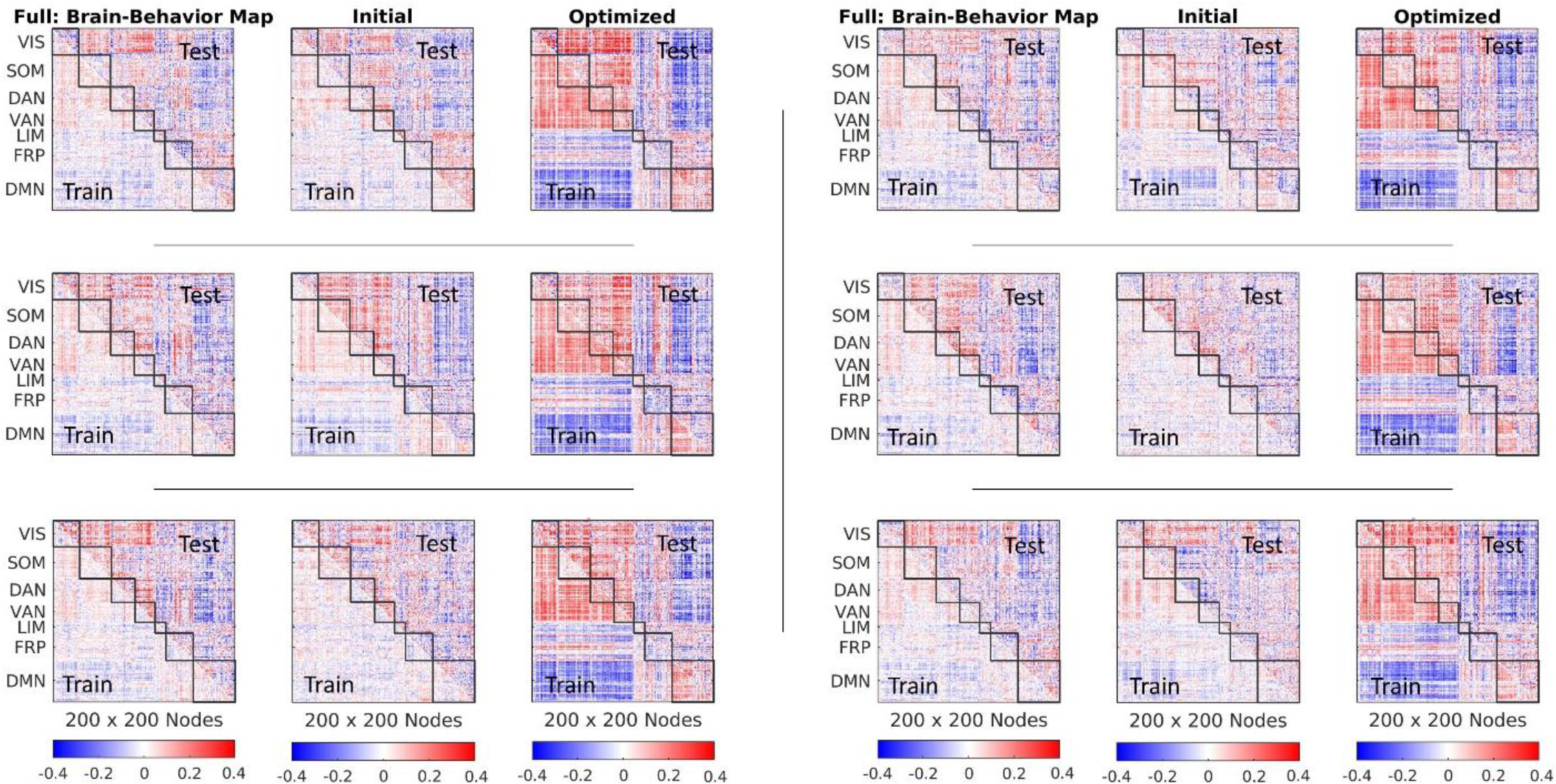
Cross validation examples of brain-behavior correlation maps for vocabulary pronunciation. Results from six cross validations with the highest train-test transfer are shown for training and testing subjects from full FC, initial template, and optimized template.

**Figure SI3.**
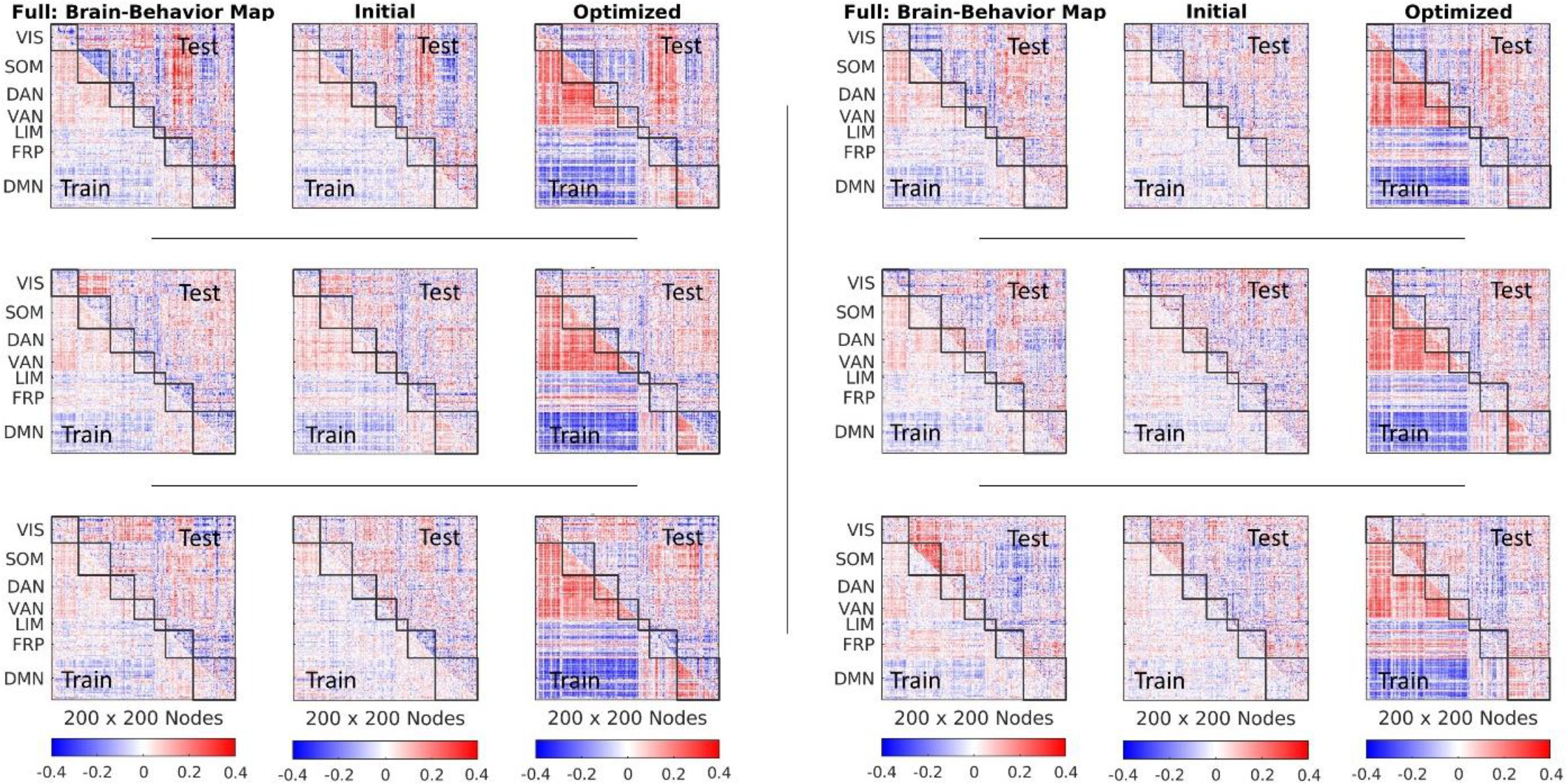
Cross validation examples of brain-behavior correlation maps for vocabulary pronunciation with inverted behavioral relations. Results from six cross validations with the lowest train-test transfer are shown for training and testing subjects from full FC, initial template, and optimized template.

**Figure SI4.**
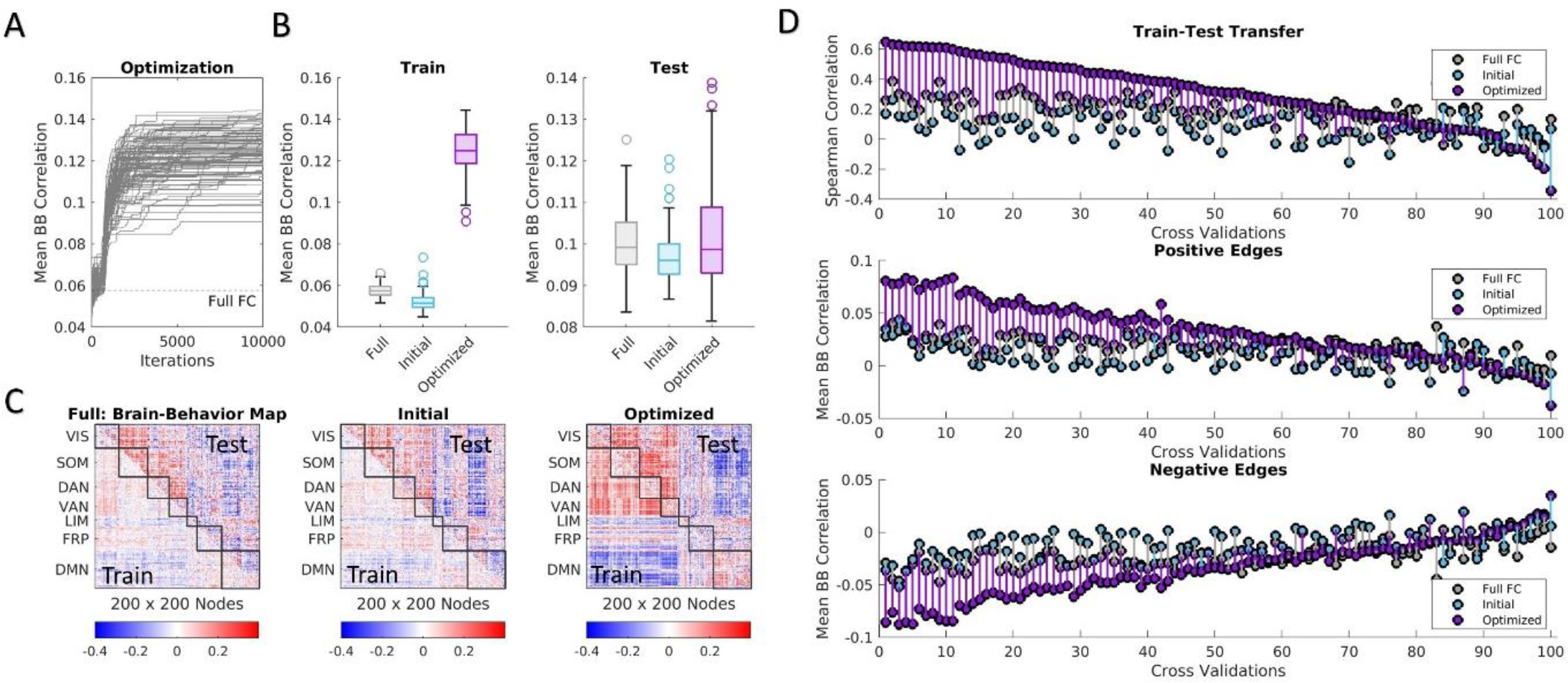
Optimization results for example behavior (vocabulary pronunciation) from filtering 5% of frames. **(A)** Mean absolute brain-behavior correlations (mean BB) across each iteration of all cross validations compared to full FC. **(B)** Performance in mean BB for brain-behavior correlation maps created from full FC, initial templates, and optimized templates for training (*left*) and testing (*right*) groups across all cross validations. **(C)** Example brain-behavior correlation maps from full FC, initial, and optimized templates from best performing cross validation for training and testing subjects. **(D)** Performance of full FC, initial, and optimized templates assessed for each cross validation using Spearman correlation between training and testing brain-behavior maps (*top*) as well as mean BB of positive edges (*middle*) and negative edges (*bottom*). Positive and negative edge results are shown for the testing group using edge masks selected from brain-behavior correlation maps of training subjects and were ordered based on performance of train-test transfer (*top*).

**Figure SI5.**
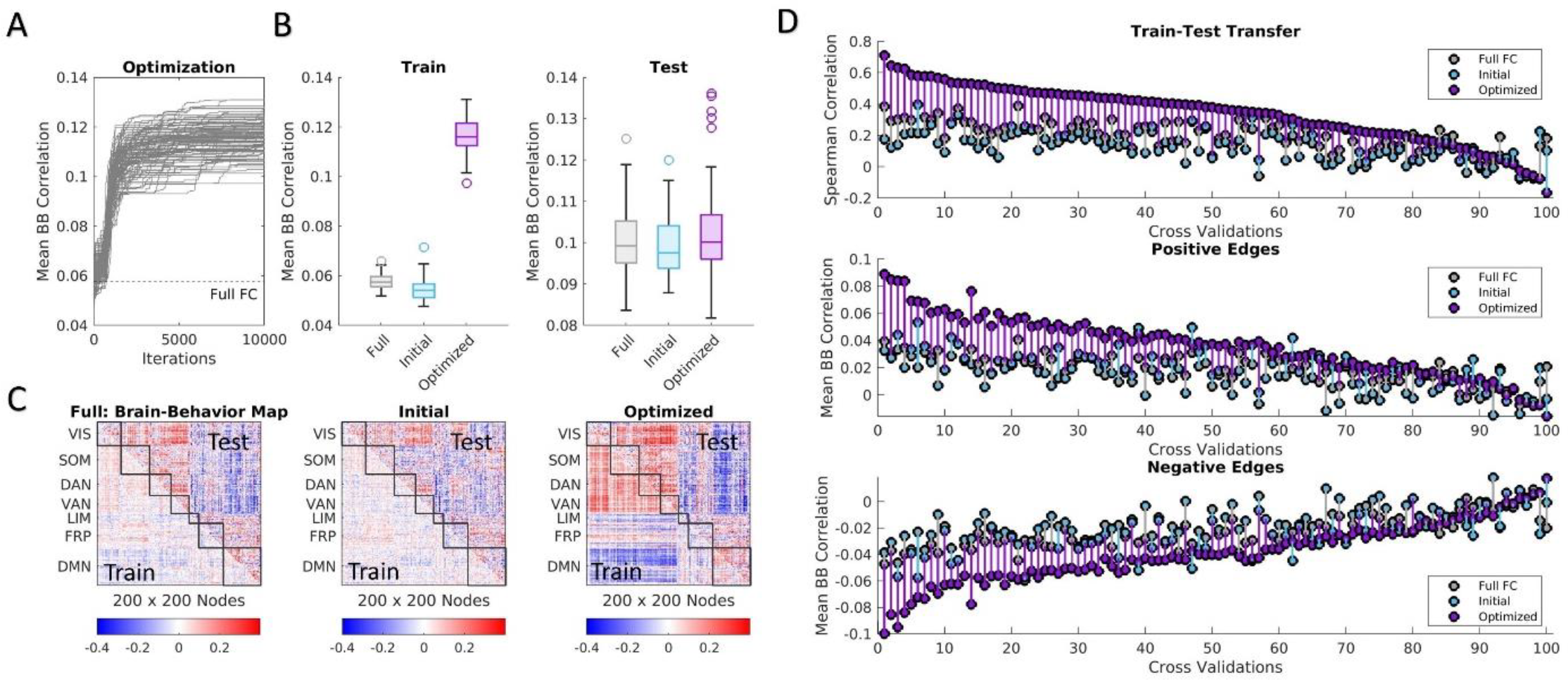
Optimization results for example behavior (vocabulary pronunciation) from filtering 15% of frames. **(A)** Mean absolute brain-behavior correlations (mean BB) across each iteration of all cross validations compared to full FC. **(B)** Performance in mean BB for brain-behavior correlation maps created from full FC, initial templates, and optimized templates for training (*left*) and testing (*right*) groups across all cross validations. **(C)** Example brain-behavior correlation maps from full FC, initial, and optimized templates from best performing cross validation for training and testing subjects. **(D)** Performance of full FC, initial, and optimized templates assessed for each cross validation using Spearman correlation between training and testing brain-behavior maps (*top*) as well as mean BB of positive edges (*middle*) and negative edges (*bottom*). Positive and negative edge results are shown for the testing group using edge masks selected from brain-behavior correlation maps of training subjects and were ordered based on performance of train-test transfer (*top*).

**Figure SI6.**
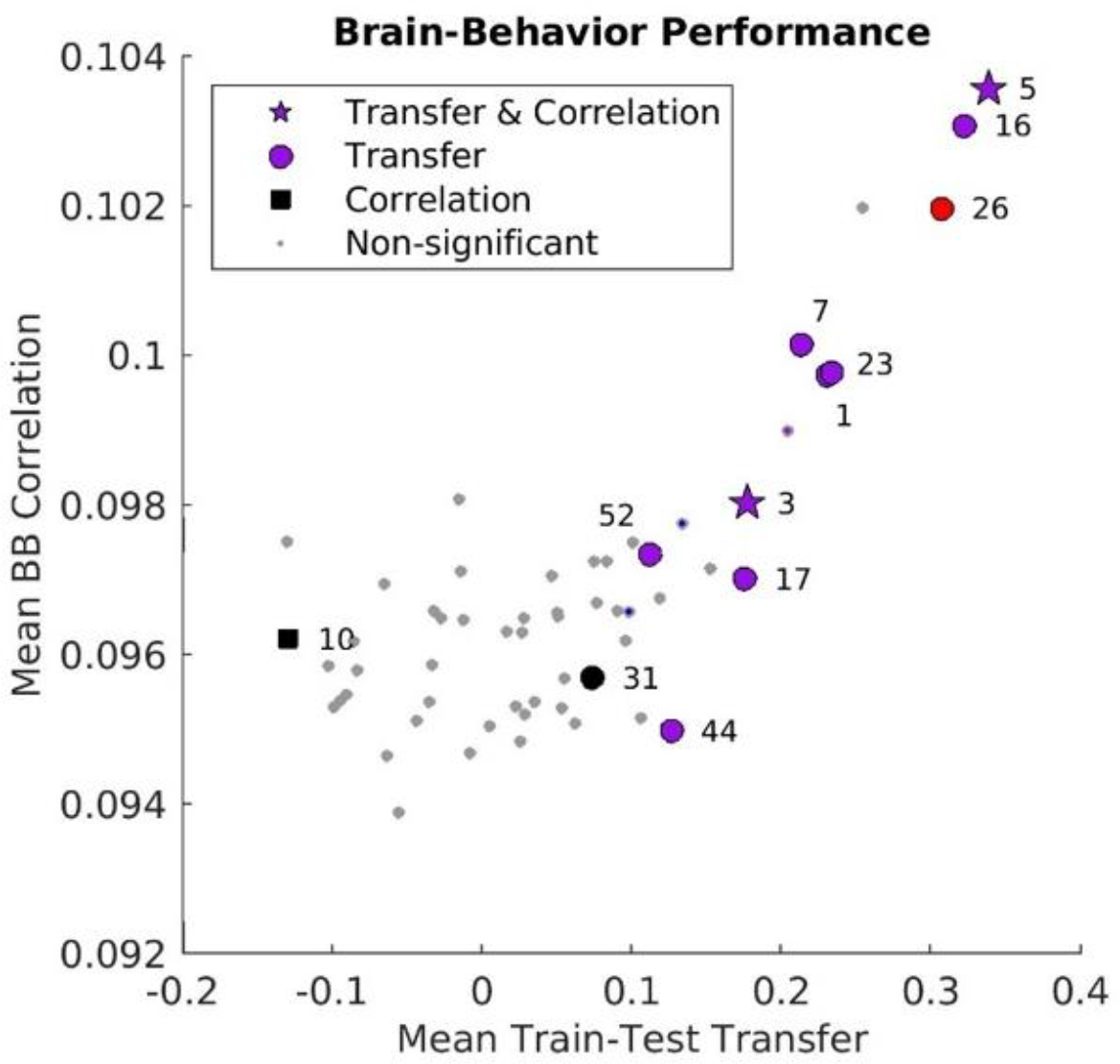
Optimization summary across all behaviors using a p < 0.001 threshold. Results from all 58 behaviors plotted as brain-behavior correlations of testing subjects versus train-test transfer (mean Spearman correlation between training and testing brain-behavior correlation maps). Across all cross validations, each behavior was compared to full FC for train-test transfer, average strength of behavioral correlations across all edges, and average strength of behavioral correlations of positive and negative edge sets. Some behaviors showed significant improvements (p < 0.001) over full FC in all mentioned categories (star), improvements in train-test transfer (circle), and average strength of all behavioral correlations (square). Behaviors labeled in purple additionally showed greater average strength of behavior correlations in both positive (red) and negative (blue) edges.

**Figure SI7.**
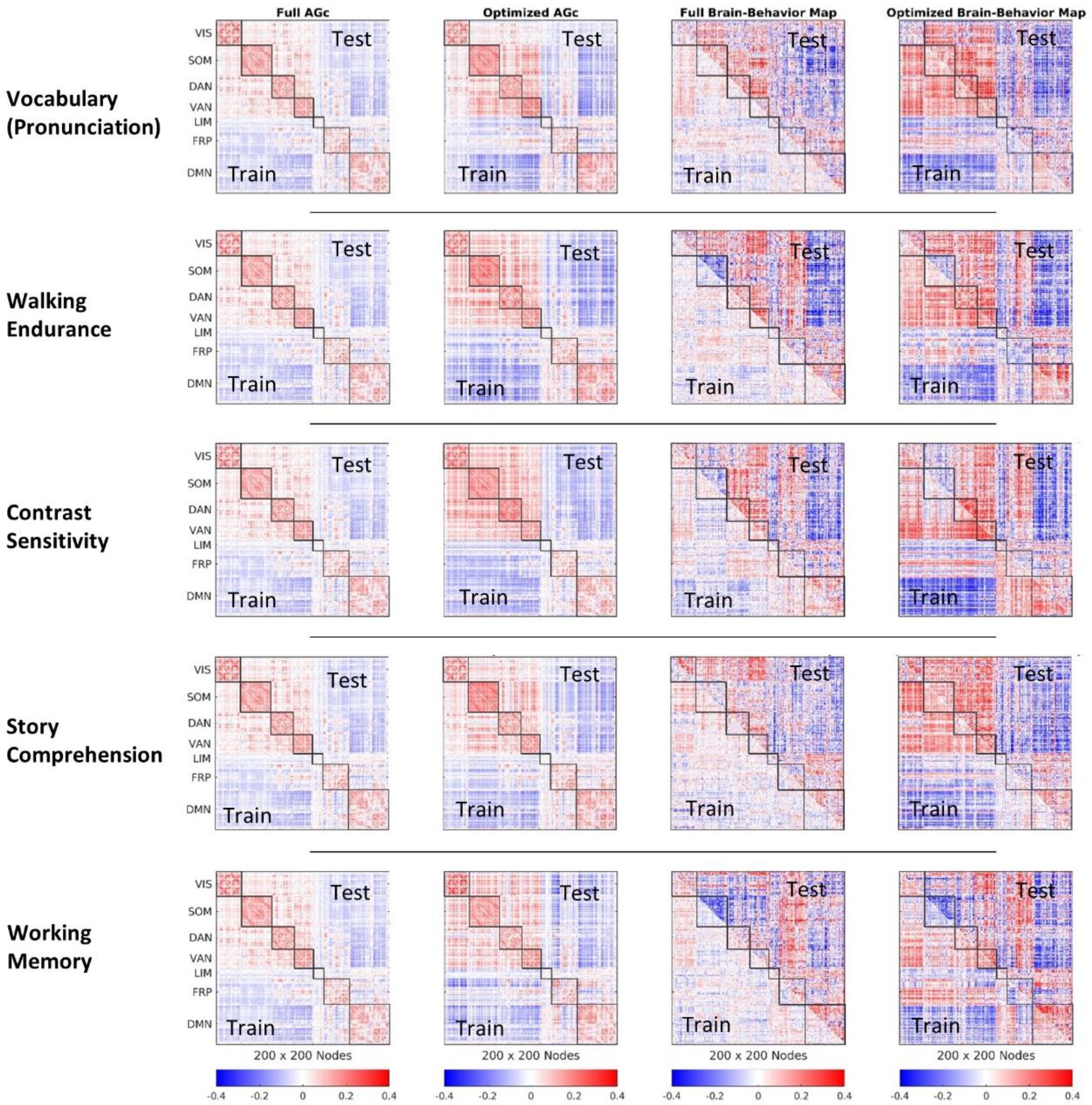
Examples of filtered AGc and corresponding brain-behavior correlation maps. Examples of subject-averaged AGcs reconstructed from filtered frames and their corresponding brain-behavior correlation maps. Results from training and testing subjects are shown for the selection of all frames (Full AGc) and from the optimized template (Optimized AGc).

**Figure SI8.**
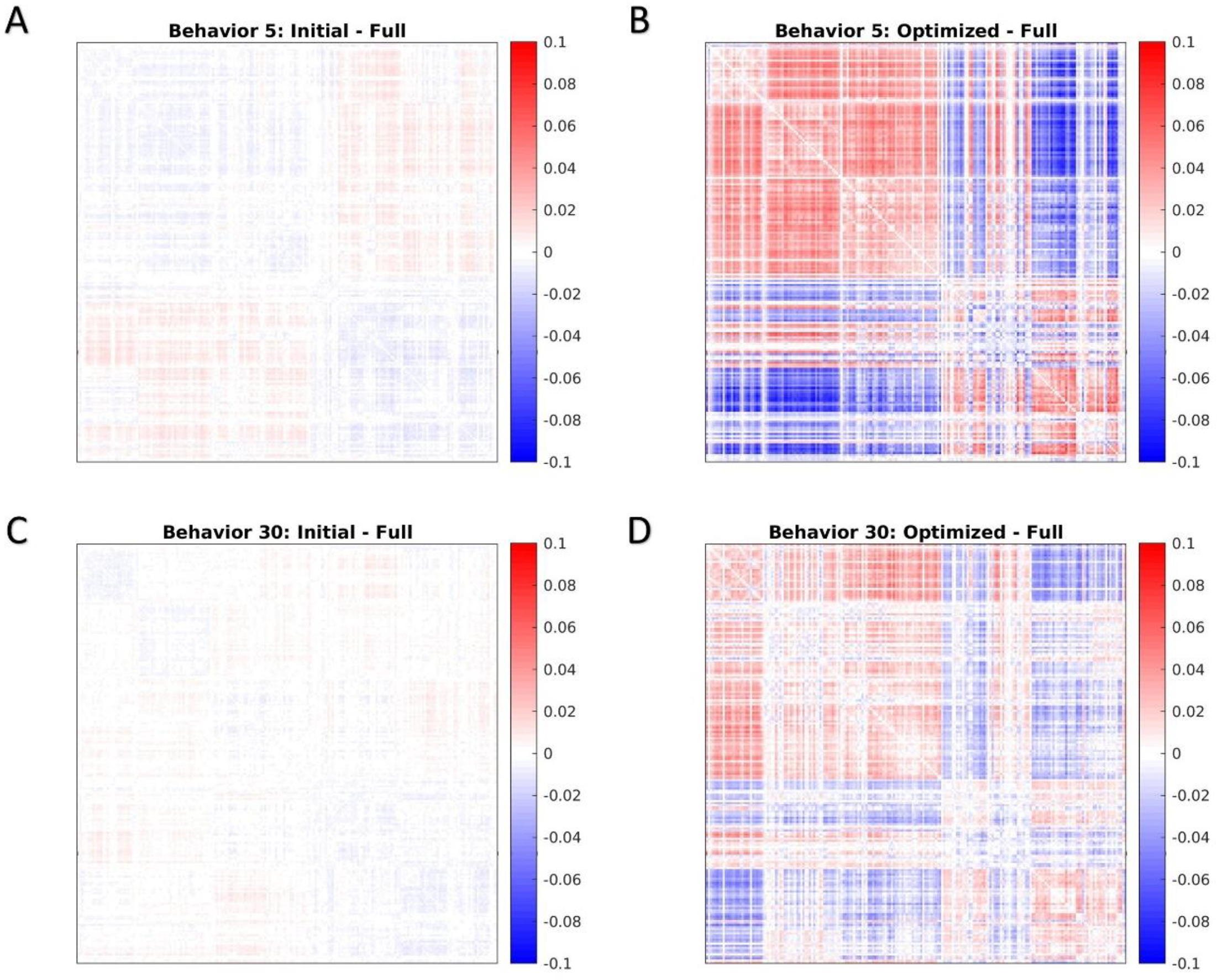
Difference between filtered AGc and full FC. **(A)** Difference between AGc from the initial template and full FC averaged across testing subjects for vocabulary pronunciation (behavior 5). **(B)** Difference between AGc from the optimized template and full FC averaged across testing subjects for vocabulary pronunciation (behavior 5). **(C, D)** Results from working memory (behavior 30) for the initial template **(C)** and optimized template **(D)**.

**Figure SI9.**
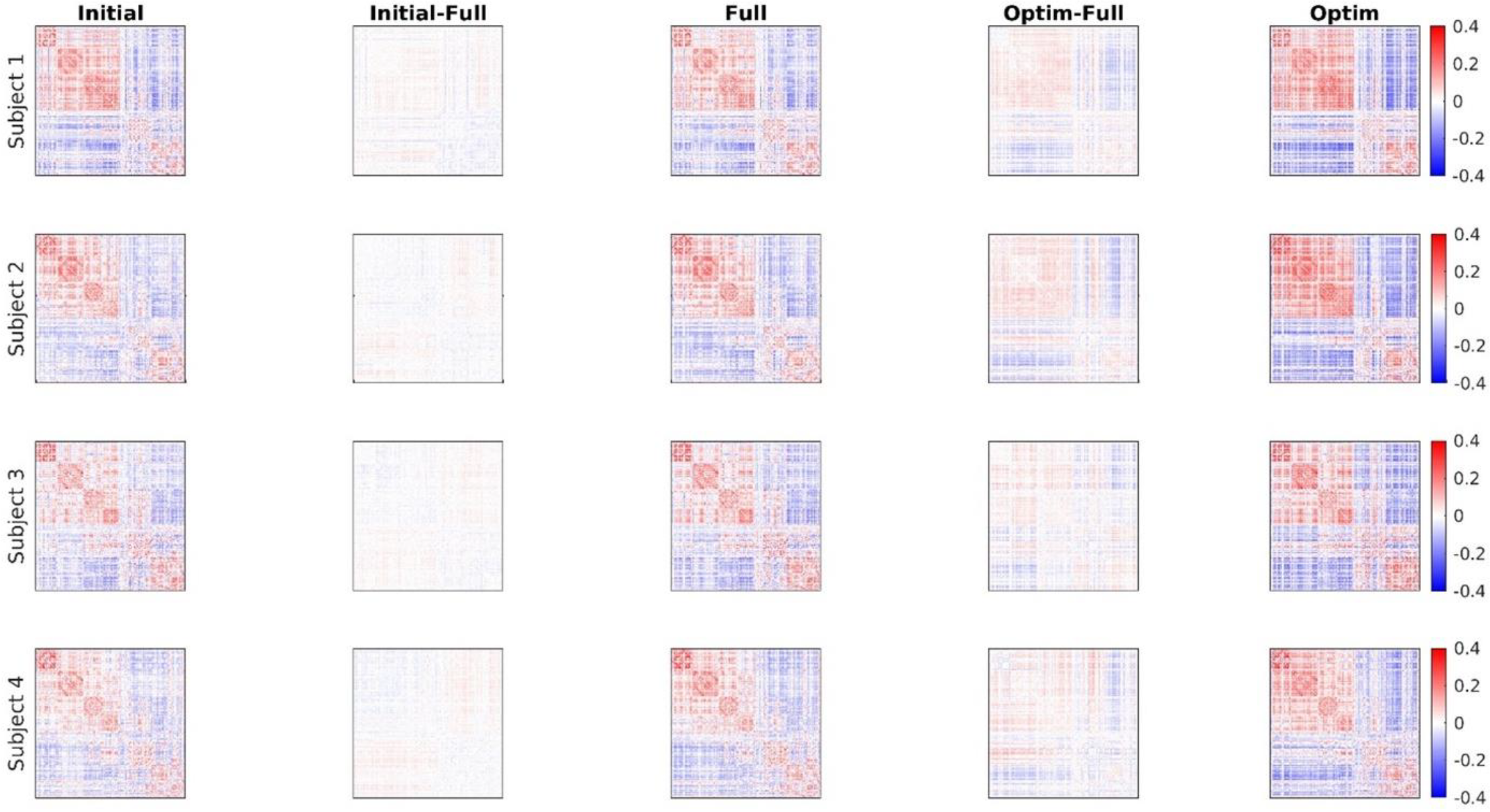
Difference between filtered AGc and full FC examples from individual subjects. Example subject AGcs are shown for frames filtered using the initial template, all frames (Full), and optimized template (Optim). Difference maps between AGc made from initial and optimized with Full are displayed separately for each example testing subject.

**Figure SI10.**
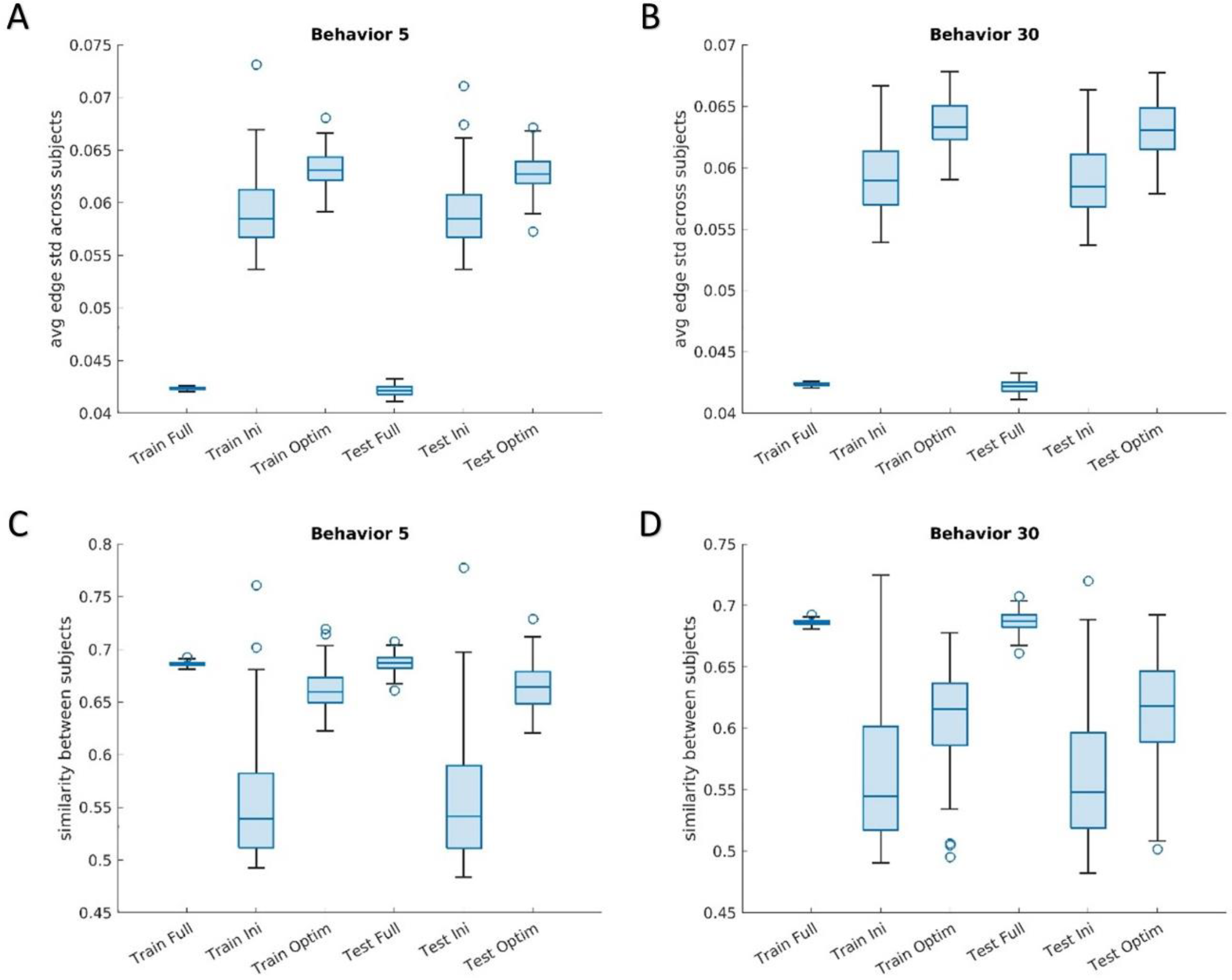
Subject variability of filtered frames. **(A, B)** Average edge standard deviation across subjects for AGc made from filtered frames using full FC, initial, and filtered templates on training and testing subject groups. **(A)** Results for vocabulary pronunciation (behavior 5) show significantly greater edge standard deviation of AGc between initial and optimized as well as full FC and optimized for both the training and testing sets (p < 10^-16^, paired-sample t-tests). **(B)** Results for working memory (behavior 30; all tests p < 10^-16^). **(C, D)** Average similarity (Spearman correlation) of AGc between subjects for full FC, initial template, and optimized in training and testing groups for vocabulary pronunciation **(C)** and working memory **(D)**. Values of subject similarity between initial and optimized as well as full FC and optimized for both the training and testing sets were significantly different for each behavior (p < 10^-10^, paired-sample t-tests).

**Figure SI11.**
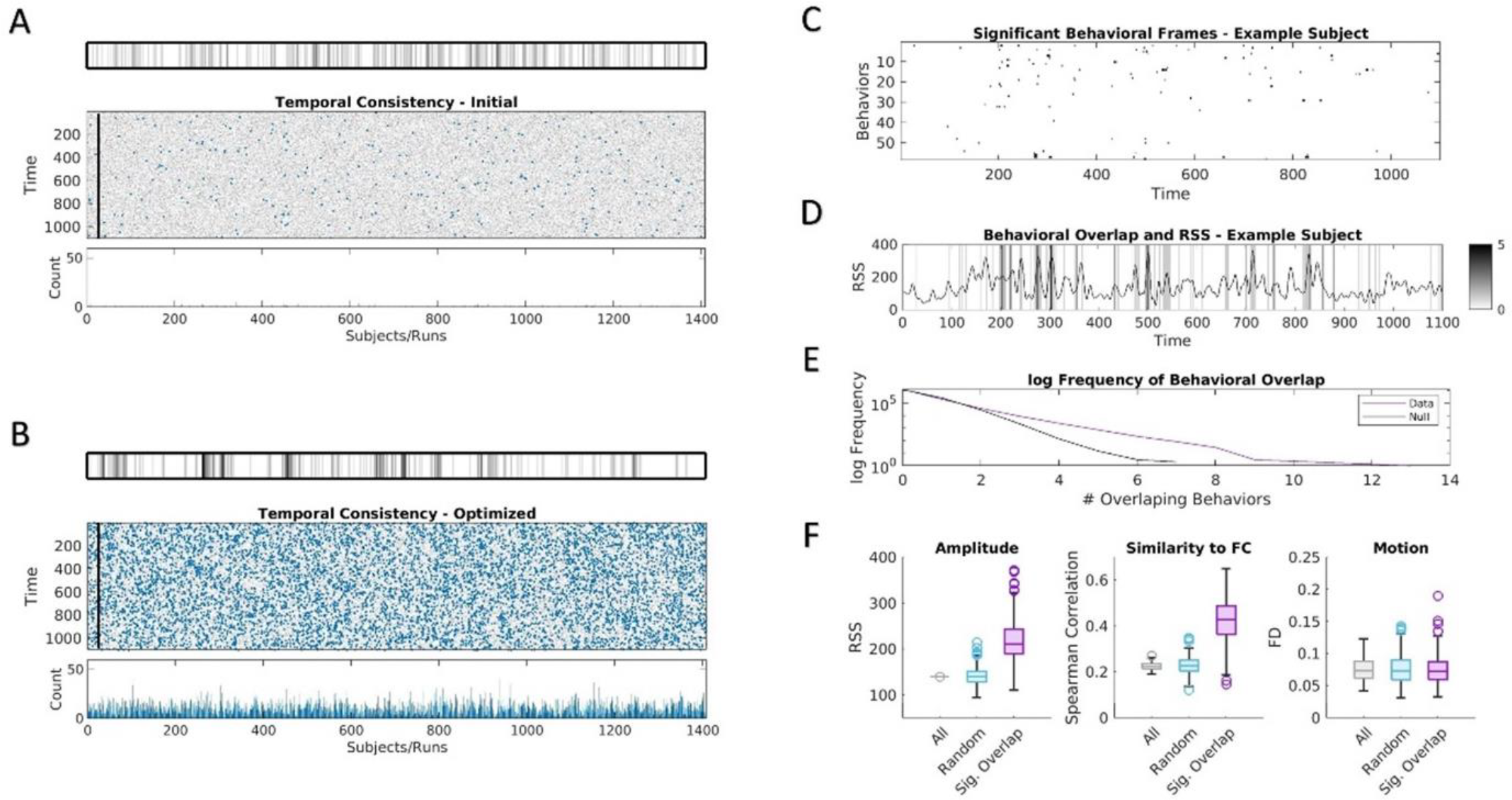
Temporal consistency and behavioral overlap using frames with significant overlap of p < 0.001. **(A)** Temporal consistency of frames filtered using the initial template of example behavior (vocabulary pronunciation) across multiple cross validations. Results are shown for cross validation training templates applied to subjects in the testing groups. Frequency of frames selected across all considered cross validations for each time point are displayed for every subject and run. Example subject shown (*top*) as plotted for every subject and run (*middle*). Time points with significant consistency between cross validations (p < 0.001; compared to 10,000 circshifted nulls) plotted in blue and the frequency of occurrence of these significant time points are shown (*bottom*). **(B)** Temporal consistency of optimized templates of example behavior (vocabulary pronunciation). **(C)** Frames with significant consistency as shown in B plotted for each behavior across time for one example subject and run. **(D)** Overlap across behaviors of significant time points as in C for the example subject and run. Corresponding RSS values displayed. **(E)** Plot of log frequency of instances of behavioral overlap of significant frames across all subjects and runs of the test group compared to distribution of 10,000 circshifted nulls. **(F)** Mean characteristics of frames across all subjects and runs for all frames, matching number of randomly selected frames, significant behavioral overlap (p < 0.001) for RSS (*left*), similarity to FC (*middle*), and motion (framewise displacement; *right*).

**Figure SI12.**
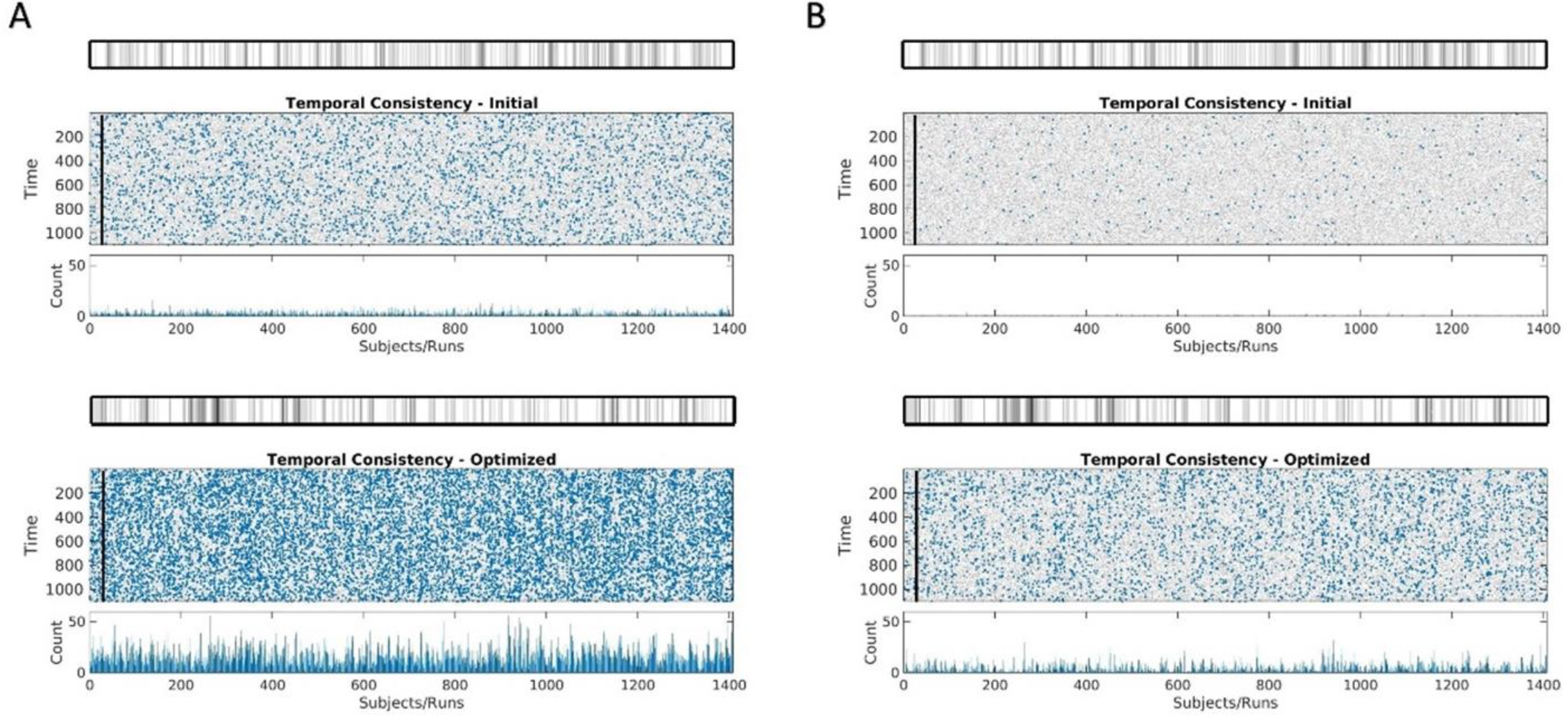
Temporal consistency and behavioral overlap of working memory. **(A)** Temporal consistency of frames filtered using initial template (*top*) and optimized template (*bottom*) using a significance threshold of p < 0.01. **(B)** Temporal consistency of frames filtered using initial template (*top*) and optimized template (*bottom*) using a significance threshold of p < 0.001.

## Notes

### Competing Interest Statement

The authors have declared no competing interest.

